# C_4_ photosynthetic pathway fluxes in transgenic rice plants

**DOI:** 10.64898/2026.05.28.728371

**Authors:** Chiara Baccolini, Stéphanie Arrivault, Florence Danila, Maria Ermakova, Hirofumi Ishihara, Regina Feil, Kavya Yalamanchili, Jane A. Langdale, Susanne von Caemmerer, Robert T. Furbank, Mark Stitt, John E. Lunn

**Author notes:** These authors contributed equally (Chiara Baccolini, Stéphanie Arrivault, Florence Danila, Maria Ermakova). Arterra Bioscience S.p.A., via Benedetto Brin, 69 80142 Naples, Italy (Chiara Baccolini). Institute of Pharmacy, Martin-Luther-University, Hoher Weg 8, 06120 Halle, Germany (Stéphanie Arrivault). School of Biological Sciences, Monash University, Melbourne, Australia (Maria Ermakova). Department of Agricultural Sciences, University of Helsinki, Helsinki, Finland (Hirofumi Ishihara). Chiara Baccolini Stéphanie Arrivault Florence Danila Maria Ermakova Kavya Yalamanchili Hirofumi Ishihara Regina Feil Jane A. Langdale Susanne von Caemmerer Robert T. Furbank Mark Stitt John E. Lunn.

## Abstract

Most land plants photosynthesize using the C_3_ pathway, in which ribulose bisphosphate carboxylase/oxygenase (Rubisco) fixes CO_2_ into 3-carbon acids. The C_4_ pathway, a biochemical CO_2_-concentrating mechanism that operates in the context of specialized leaf anatomy to concentrate CO_2_ around Rubisco, is more efficient. Introduction of the C_4_ pathway into the C_3_ crop rice could increase yield by 50%. Expression of five C_4_ enzymes in transgenic rice previously led to flux through the first step. However, there was no evidence for flux later in the cycle. Here we developed new transgenic rice lines and novel protocols to detect C_4_ cycle activity: CO_2_ fixation into C_4_ acids by carboxylation of a C_3_ compound, decarboxylation, refixation of CO_2_ by Rubisco, and regeneration of the C_3_ donor. We demonstrate that these four core C_4_ reactions are operating in rice, establishing the *in vivo* flux framework needed to progress towards a functional carbon-concentrating mechanism.

## INTRODUCTION

Photosynthesis is the process by which almost all carbon (C) enters the biosphere, including all food and feed production in agriculture. There is a pressing need to increase agricultural yield to keep pace with the growing world population and to stabilise yield in the face of climate change ^1^. Most terrestrial plants carry out C_3_ photosynthesis. After entering the leaf, CO_2_ is assimilated by ribulose-1,5-bis-phosphate carboxylase/oxygenase (Rubisco) in the Calvin-Benson cycle (CBC) to produce the 3-carbon metabolite 3-phosphoglycerate (3PGA). Rubisco also catalyses a competing reaction with O_2_, producing 2-phosphoglycolate, an inhibitory metabolite that is recycled by a process called photorespiration. This infidelity consumes up to 50% of the ATP and NADPH generated in the light reactions and imposes constraints on stomatal conductance and nitrogen allocation ^2,3^. Biochemical and morphological adaptations to concentrate CO_2_ around Rubisco and suppress the side-reaction with O_2_ have emerged repeatedly during evolution; in cyanobacteria, green algae and terrestrial plants. One of these adaptations is C_4_ photosynthesis ^4-6^, which evolved independently in over 60 plant lineages over the last 30 million years ^3^. C_4_ plants initially fix CO_2_ into C_4_ acids in mesophyll cells using the O_2_ insensitive phospho-*enol-*pyruvate carboxylase (PEPC). The C_4_ acids then diffuse to neighbouring bundle sheath cells where they are decarboxylated to release CO_2_ for refixation by Rubisco (Figure 1). This CO_2_-carbon concentrating mechanism (CCM) leads to higher rates of photosynthesis and better water and nitrogen use efficiency than found in C_3_ plants ^2,7^. Installation of a C_4_ pathway into staple crop plants like rice (*Oryza sativa*) that operate C_3_ photosynthesis is predicted to increase yields by up to 50% ^8^.

**Figure 1.**
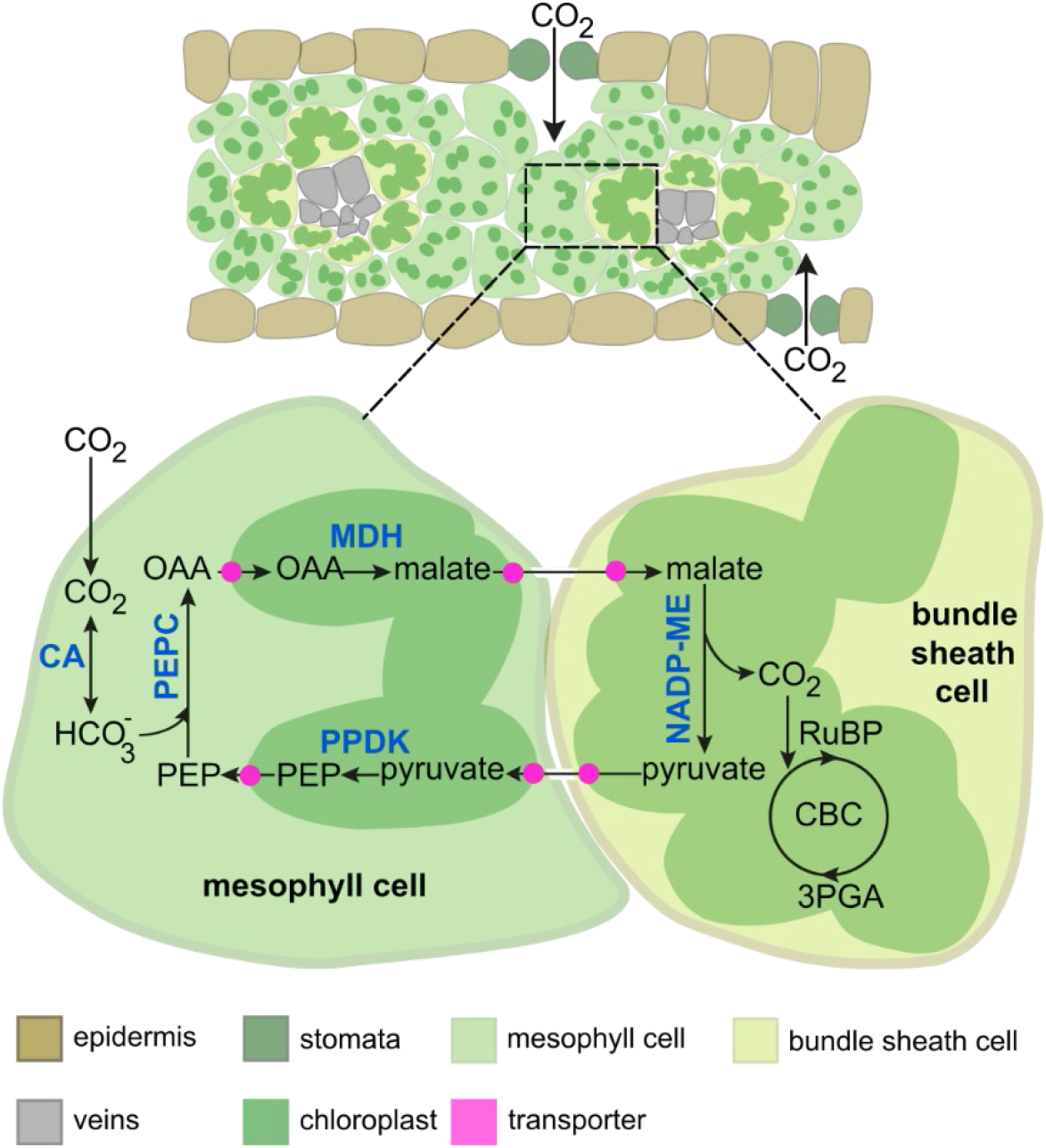
Schematic of C_4_ anatomy and metabolism. Transverse sections of C_4_ leaves reveal veins that are encircled by large bundle sheath (BS) cells that are themselves surrounded by mesophyll (M) cells. In this arrangement, each pair of veins is separated by two BS and two M cells. C_4_ metabolism is shared between these two cell-types. In the M cells, CO_2_ is interconverted with bicarbonate (HCO_3_) by carbonic anhydrase (CA) and bicarbonate is assimilated by phosphoenolpyruvate carboxylase (PEPC) to produce the C_4_ acid oxaloacetate (OAA). In the majority of C_4_ grasses ^54^, OAA is transported into the chloroplast where it is reduced by malate dehydrogenase (MDH) to malate. Malate then exits the chloroplast and diffuses through plasmodesmata into the BS cells where it is decarboxylated by NADP-dependent malic enzyme (NADP-ME) in the chloroplasts (as shown) or by NAD-ME in the mitochondria, and/or by PEP carboxykinase in the cytoplasm ^55^. The released CO₂ is then re-fixed by Rubisco in the Calvin-Benson cycle (CBC). The remaining three-carbon compound, pyruvate, diffuses back to the M cells, where it is transported into the chloroplast and phosphorylated by pyruvate phosphate dikinase (PPDK) to regenerate the initial CO₂ acceptor, phosphoenolpyruvate (PEP). RuBP = Ribulose 1,5-bisphosphate; 3PGA = 3-Phosphoglycerate.

Establishment of a C_4_ cycle in C_3_ plants such as rice requires the modification of both anatomical and biochemical traits. Specifically, chloroplasts need to be functionalized in the bundle sheath cells that surround the leaf veins ^3,9,10^, the spacing between veins needs to be reduced to bring the ratio of bundle sheath to mesophyll cells close to one i.e., the number of mesophyll cells between veins needs to reduce from ∼6 to 9 to only 2 ^11,12^, and the individual C_4_ enzymes need to accumulate and be active at high levels specifically in either the mesophyll or bundle sheath cells. Several studies that have engineered genes encoding individual C_4_ enzymes into C_3_ plants have reported enhanced extractable enzyme activity but no enhancement of *in vivo* enzyme activity ^13-16^. When the C_4_ pathway was discovered, information about *in-vivo* fluxes was provided by ^14^CO_2_ pulse-chase labelling experiments. These revealed strong labelling of C_4_-acids in the pulse, and rapid loss of label from C_4_-acids and continued movement of label into 3PGA in the chase ^4,5^. In our previous attempts to engineer a C_4_ cycle into C_3_ plants, Lin *et al*. (2020)^17^ reported the introduction of genes encoding a minimum set of four enzymes necessary for the C_4_ cycle in maize (*ZmPEPC*, *ZmNADP-MALATE DEHYDROGENASE (MDH)*, *ZmNADP-MALIC ENZYME (ME)*, *ZmPYRUVATE ORTHOPHOSPHATE diKINASE (PPDK*; see Figure 1) into *O. sativa indic*a cv. IR64, and Ermakova *et al*. (2021)^18^ reported introduction of this minimum set plus carbonic anhydrase (CA) into *O. sativa japonica* cv. Kitaake. Suitable promotors were employed to drive high levels of transgene expression in the correct cell type and in both studies extractable enzyme activities were higher than in the corresponding wild-type controls. Notably, a 3-fold increase in extractable PEPC activity led to a 10-fold increase in labelling of the 4 carbon compounds malate and aspartate in ^13^CO_2_ pulse experiments, suggesting that the carboxylation step of the C_4_ cycle was functioning in the transgenic lines ^18^. However, ^13^CO_2_ pulse-chase experiments failed to find evidence for enhanced decarboxylation despite a 7-fold increase in extractable NADP-ME activity ^18^ and no direct way was found to measure flux at PPDK. Thus, whilst introduction of a minimal set of C_4_ cycle enzymes led to enhanced PEP carboxylation, there was no evidence for enhanced flux at later steps in the C_4_ cycle.

Here, we report new transgenic rice lines with all 5 enzymes of the C_4_ pathway appropriately expressed, including lines where NADP-ME expression is bundle sheath cell-specific and extractable activity is over half of that in C_4_ plants. We also report the development of new protocols that are sufficiently sensitive to detect small increments in flux at the four core steps of the C_4_ cycle: carboxylation by PEPC, decarboxylation of C_4_ acids, refixation of the released CO_2_, and regeneration of PEP from pyruvate by PPDK. Characterization of the transgenic rice lines revealed flux through each step of the C_4_ cycle, a landmark achievement in the pursuit of C_4_ rice.

## RESULTS

### Generation of transgenic rice lines accumulating high levels of all five core C_4_ enzymes

To generate transgenic rice lines expressing all five of the core C_4_ enzymes from maize at high levels, we sought to elevate expression of *ZmNADP-ME*, the transgene that was only weakly expressed in previously published lines ^18^. To this end we used a synthetic transcription factor (dTALE) driven by a bundle sheath (BS) cell-preferential promoter together with cognate promoters (STAP) to drive expression of the maize *NADP-ME* gene (^19^, Figure S1A). T1 lines were identified that accumulated NADP-ME protein up to levels around half of that detected in maize (Figure S1B), one of which accumulated ME specifically in BS cells (3.4_ME_BS_) (Figure S1C). In lines generated with a second dTALE-STAP construct, NADP-ME protein was similarly detected at high levels but despite the presence of the BS-preferential promoter in the transgene construct, protein was detected in both BS and mesophyll (M) cells (Figure S1C), presumably as a consequence of genomic insertion site (Figure S1A-B). Both lines (henceforth referred to as ‘high ME lines’) (Figure S1D) were crossed into two established transgenic rice cv. Kitaake lines (29 and B6) that had substantially increased protein abundance and/or activities of CA, PEPC, NADP-MDH and PPDK in M cells ^18^, to generate ME_BS_ x 29 (29A & 29C) and ME_BS&M_ x B6 (B6A & B6B) F1 plants. F1 plants were screened for the presence of both transgene constructs and for the accumulation of all five C_4_ proteins before being selfed to generate F2 lines (Figure 2A). In homozygous F2 plants generated from the F1 lines, NADP-ME protein accumulated to levels over 25% of those seen in maize (Figure 2B) and *in vitro* NADP-ME activities were 12-16% (ME_BS&M_ x B6) and 30-38% (ME_BS_ x 29) of those in the C_4_ species *Setaria viridis* and *Sorghum bicolor* respectively (Figure 2C; Table S1). As in the parental 29 and B6 lines, immunoblotting of CA, PEPC, NADP-MDH and PPDK proteins indicated levels ranging from 20% to >100% of those in maize (Figure 2B) and accumulation preferentially in M cells (Figure 2D). Like the parental high NADP-ME lines, ME_BS_ x 29 lines accumulated NADP-ME specifically in BS cells whereas ME_BS&M_ x B6 lines accumulated NADP-ME in both BS and M cells (Figure 2E). Given that low activity of endogenous transporters may preclude movement of malate and aspartate into the BS chloroplasts and/or that diffusion of C_4_ acids from M to BS cells may be compromised by the C_3_ leaf anatomy, the ME_BS&M_ x B6 lines provided a biological chassis in which to test whether decarboxylation of C_4_ acids generated by *Zm*PEPC could occur within the M cells of transgenic rice lines. More importantly, the ME_BS_ x 29 lines provided a chassis to test for compartmentalized C_4_ cycle function between M and BS cells. Overall, *in vitro* extractable activity of the C_4_ enzymes PEPC, NADP-ME and PPDK in the parental and derived transgenic rice lines were about 3.4%, 12-38% and 15% of those in maize (Table S1).

**Figure 2.**
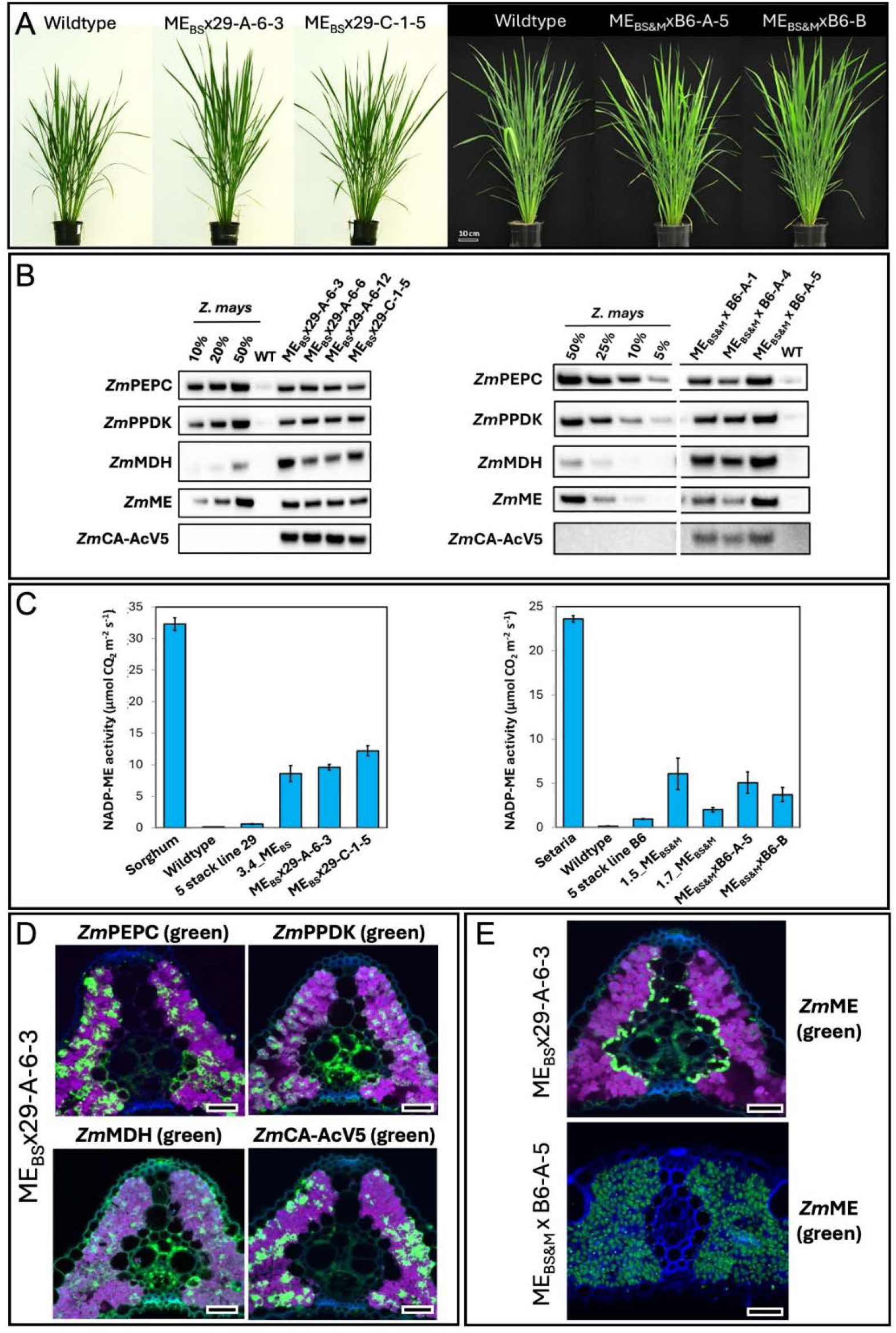
Accumulation of C_4_ proteins in transgenic rice lines. **A)** Normal growth phenotype of F2 transgenic plants at mid-tillering stage. **B)** Western blot showing accumulation of the five maize C_4_ maize proteins (PEPC, PPDK, MDH, ME and CA) in F2 lines. **C)** Activity of NADP-ME in parental lines (29/B6 & 3.4_ME_BS_/1.5_ME_BS&M_/1.7_ME_BS&M_) and in F2 lines relative to C_4_ sorghum, C_4_ setaria and wildtype rice, n = 3 to 5. NADP-ME activities are shown on a leaf area basis to compare with gas exchange flux. Table S1 shows enzyme activities on a fresh weight basis, and compared to those reported in maize. **D)** Immunolocalization of the four mesophyll-specific C_4_ maize proteins in mesophyll cells of one of the selected F2 lines (ME_BS_x29-A-6-3). Images are pseudo-coloured: green-protein of interest, magenta-chlorophyll autofluorescence, blue-cell wall. Scale bars = 20 µm. **E)** Immunolocalization of maize NADP-ME protein in BS chloroplasts of F2 line ME_BS_x29-A-6-3 and in both BS and M cells of F2 line ME_BS&M_xB6-A-5. Supported by Figure S1 and Table S1.

### Up to ten-fold enhancement of CO_2_ fixation into C_4_ acids in transgenic rice lines

To determine whether all steps of the C_4_ cycle functioned in the newly generated rice lines, we first assessed whether PEPC-mediated carboxylation occurred at similar levels in the ME_BS_ x 29 and ME_BS&M_ x B6 lines as in the published parental lines ^18^. In a C_4_ plant, following interconversion of CO_2_ and bicarbonate by CA, assimilation of bicarbonate into malate and/or aspartate by PEPC is easily detected by supplying a short pulse of ^14^CO_2_ or ^13^CO_2_ and investigating label incorporation into the major downstream products of the PEPC reaction – the C_4_ acids malate and aspartate (Figure 3A). However, this analysis is not suitable to search for low-level activity in transgenic rice lines against a background of rapid C_3_ photosynthesis because the initial labelling kinetics of malate and aspartate include not only incorporation of ^13^C from bicarbonate into the C4 position, but also incorporation of label that moves from the CBC into positions C1-C3 of the C_4_-acids ^20^ (Figure 3A). To circumvent this problem, we calculated the rate of carboxylation by summing the abundance of the labelled isotopologues of malate and aspartate (i.e., m*_1_*, m*_2_*, m*_3_* + m*_4_*, nmol g^-1^ FW) (see Supplementary Calculations). This analysis was applied to three ^13^CO_2_ pulse-labelling datasets: one with the parental lines only (published in ^18^) and two new datasets generated with the ME_BS_ x 29 and ME_BS&M_ x B6 lines that used very short, precise pulses of ^13^CO_2_ (420 ppm) and chases with unlabelled CO_2_ (420 ppm). Notably, all of the transgenic lines exhibited a faster rise in the abundance of the summed labelled isotopologues of malate (Figure 3B, Figure S2, Supplementary Datasets 1A-C). There was also rapid labelling of aspartate, suggesting rapid equilibration of oxaloacetate, malate and aspartate by either the endogenous or introduced MDH and endogenous aspartate aminotransferase (AAT). Importantly, the summed abundance of labelled malate and aspartate isotopologues provided an initial estimate of flux at PEPC with an enhancement of between 4- to 10-fold relative to wild-type observed in all of the new transgenic lines (Figure 3B and legend), consistent with previous observations in the parental 29 and B6 lines ^18^.

**Figure 3.**
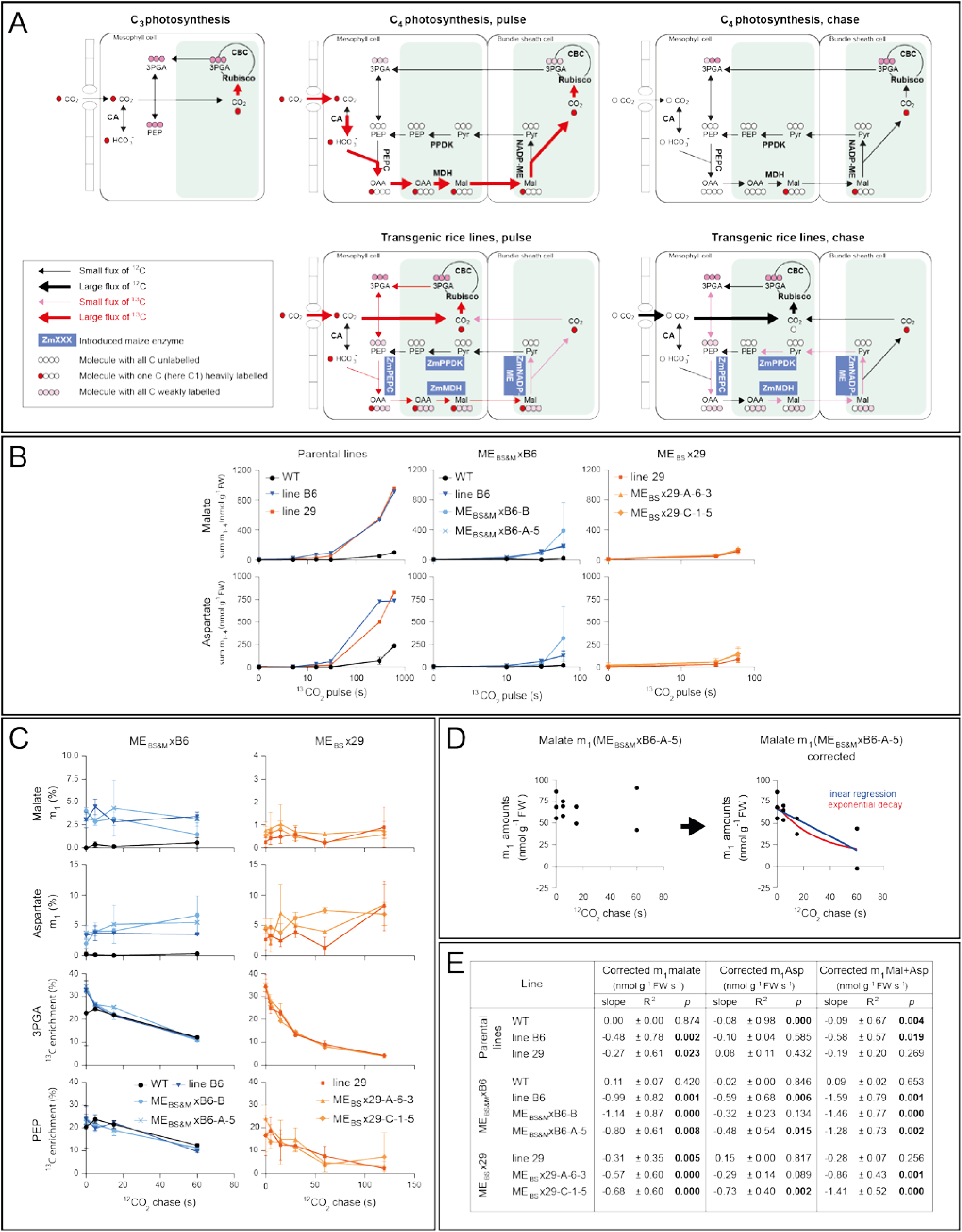
Accumulation and decay of labelled isotopologues of malate and aspartate during pulse-chase labelling with ^13^CO_2_. **A)** Schematics to show flow of label in C_3_ photosynthesis, C_4_ photosynthesis, and in transgenic rice lines. In the pulse PEPC flux is measured by summing labelled malate and aspartate molecules. In the chase, decarboxylation flux is measured as the loss of labelled malate and aspartate molecules, after correcting for molecules that are synthesised from labelled PEP during the chase. The displays give a schematic overview of the flow of ^13^C and ^12^C in the labelling experiments including the intramolecular labelling patterns in the 4-carbon and 3-carbon intermediates in the C_4_ cycle, the CBC and in PEP. Note that in C_4_ photosynthesis there is a large delay before label moves from 3PGA into PEP and, initially, label incorporation by PEPC is from [^13^C]HCO_3_ into the C4 position of C_4_ acids ^48,56^. In rice and in transgenic rice lines expressing low amounts of C_4_ enzymes in a basically C_3_ background, label moves rapidly from 3PGA into PEP, leading to PEPC also incorporating label from PEP into positions C1-C3. Malate is shown as an example for a C_4_ acid. It is the main C_4_ acid in maize and is the substrate in the transgenic rice lines for the introduced ZmNADP-ME. The OAA produced by PEPC in the M cells is converted to malate via NADP-malate dehydrogenase. Combined action of malate dehydrogenase and aspartate amino transferase also facilitates rapid movement of label between malate, OAA and aspartate both in the M cells and the BS cells and, like malate, aspartate moves from M cells to BS cells (not shown). As interconversion of malate and aspartate occurs without rearrangement of label between the positions in the C_4_ acids, the intramolecular labelling pattern in aspartate is like that of malate. **B)** Plots showing the summed amounts (nmol g^-1^FW) of the m*_1_*, m*_2_*, m*_3_*, and m*_4_* isotopologues (where *_n_*is the number of ^13^C atoms incorporated in the metabolite) of malate (upper panel) and aspartate (lower panel) during pulse labelling in three different experiments. Every PEPC reaction in the pulse leads to the formation of a labelled isotopologue. The initial slope (nmol g^-1^FW s^-1^) is a first approximation for flux at PEPC. Summing of labelled isotopologues rather than total ^13^C incorporation excludes ^13^C that is fixed by Rubisco and moves via 3PGA into PEP, leading to appearance of label in the C1-C3 positions of C_4_ acids. Accumulation rates (nmol g^-1^ FW s^-1^) were estimated by linear regression on initial time points as 0.48-0.52 in two experiments with wild-type, compared to 4-8-5.1 in two experiments with paternal line B6, 5.3 and 4.7 in ME_BS&M_ x B6-A-5 and ME_BS&M_ x B6-B, 1.9-3.6 in two experiments with paternal line 29 and 4.8 and 5.1 in ME_BS_ x 29-A-6-3 and ME_BS_ x 29-C-1-5. The plots show mean ± SD, n = 1 to 4, 2 to 3 and 3 to 7 for each labelling experiment. See Figure S2 for labelling kinetics of all isotopologues of these and further metabolites, Supplementary Datasets 1A-C for raw data and Supplementary Calculations. **C)** Fractional abundance of m_1_ malate and m*_1_*aspartate (% m*_1_* malate, % m*_1_* aspartate) and % enrichment in 3PGA and PEP in the chase in the B6 and 29 parental lines plus ME_BS&M_ x B6 and ME_BS_ x 29 lines. Line B6 has 7-fold enhanced NADP-ME activity, ME_BS&M_ x B6-A-5 has 35-fold (in both BS and M cells), line 29 has 4-fold and lines ME_BS_ x 29 A-6-3 and C-1-5 have up to 90-fold enhanced NADP-ME activity specifically in BS cells (Figure 1C, Table S1, Figure S1). See Figure S3 for labelling kinetics of all isotopologues of these and further metabolites and Supplementary Datasets 1A-C for raw data. **D)** Correction of amount of malate m*_1_* to exclude m*_1_* malate that is synthesized in the chase from m*_1_* PEP. Label incorporated into 3PGA and other CBC intermediates in the preceding pulse continues to flow via PEP into positions C1-C3 of malate and aspartate in a chase (see panel A). This will result in continued formation of m*_1_*, m*_2_* and m*_3_* malate and aspartate in the chase, that can mask any loss of ^13^C from position C4 of C_4_ acids due to decarboxylation. The rate of, for example, m*_1_* malate formation in a given time interval can be estimated as the average m*_1_* abundance in PEP in that time interval multiplied by the rate of synthesis of malate by PEPC. As an example, the raw and corrected data are shown for line ME_BS&M_ x B6-A-5; the display shows all individual samples before and after correction. The decay of the corrected m*_1_* abundance provides evidence for decarboxylation in the chase. The corrected data are fitted with a linear regression (blue line) and an exponential decay curve (red line). See Figure S4A & B for analogous plots of all other genotypes and Supplementary Calculations for raw data. **E)** Summary of the slopes, R_2_ and p-values for the decay of corrected m*_1_* malate abundance and of corrected m*_1_* aspartate abundance estimated by linear regression. The slope provides an estimate of the rate of ^13^C loss from the C_4_ acid (nmol ^13^CO_2_ g FW^-1^ s^-1^). Regression analyses with confidence interval 95% were performed using Excel Data Analysis ToolPack (version 2408),with the the null hypothesis states that the slope coefficient is zero. The actual rate of decarboxylation will be higher because only part of the C_4_ acid pool is labelled after the preceding 15-30 s pulse. Values estimated by fitting an exponential decay curve are given in Figure S4C. Abbreviations: Mal, malate; Asp, aspartate. Supported by Figures S2-S4 and Supplementary Datasets 1A-C.

### Decarboxylation of C_4_-acids is detected by pulse-chase in transgenic rice lines

Having established enhanced flux through PEPC in transgenic rice lines following a ^13^CO_2_ pulse, we next sought to determine whether the fixed C_4_ acids were subsequently decarboxylated during a ^12^CO_2_ chase (Figure 3A). Decarboxylation was not detected in previous experiments with the parental 29 and B6 lines which have up to a 7-fold increase in extractable NADP-ME activity compared to WT ^18^ but the new lines generated here have up to 20 times more extractable NADP-ME activity than the previously published lines (Figure 2C, Table S1). We therefore looked for a decrease of label in malate or aspartate, or an increase of label in 3PGA following a ^13^CO_2_-pulse/^12^CO_2_-chase in the ME_BS_ x 29 and ME_BS&M_ x B6 lines. Surprisingly, however, no increased flux was detected (Figure 3C, Figure S3, Supplementary Datasets 1A-C). We thus hypothesized that the standard analysis of labelling kinetics, as performed in ^18^, is unsuitable for detecting enhanced decarboxylation against a background of rapid C_3_ photosynthesis. This is because in C_3_ plants, labelling of 3PGA is dominated by direct Rubisco assimilation of both ^13^CO_2_ in the pulse and ^12^CO_2_ in the chase, and because the slow decay of PEP labelling during the chase leads to continued incorporation of ^13^C into the C1-C3 positions of malate and aspartate (Figure 3A), masking any loss of ^13^C due to decarboxylation of the C4 position.

To assess the extent to which background C_3_ photosynthesis was preventing detection of decarboxylation in the transgenic rice lines, we estimated the rate of synthesis of malate and aspartate from labelled PEP in the chase. During the chase in unlabelled CO_2_, sequential action of PEPC, NADP-MDH and AAT will convert m*_1_*isotopologues of PEP into m*_1_* isotopologues of malate and aspartate. Using the pulse-chase datasets from the parental lines ^18^ and the ME_BS_ x 29 and ME_BS&M_ x B6 lines, we therefore estimated how much m*_1_*malate and m*_1_* aspartate would be produced from m*_1_*PEP at different times in the chase. This was subtracted from the m*_1_*malate and m*_1_* aspartate at that time to estimate how much of the m*_1_* malate and m*_1_* aspartate that was produced in the pulse was left at any given time in the chase (see Supplementary Calculations). As an example, Figure 3D shows the measured and corrected abundance of m*_1_* malate during the chase for one transgenic line, with the slope providing an estimate of the rate of release of ^13^CO_2_ by decarboxylation. A minimum slope was estimated by linear regression (Figure 3D) and a further estimate was obtained by fitting an exponential decay curve to each set of data points and calculating the initial slope at the beginning of the chase (Figure S4, Supplementary Calculations). Somewhat higher rates of decarboxylation were obtained by fitting an exponential decay curve but as noise was often higher, we focused on the conservative estimates provided by linear regression. Analogous corrections were performed for all experiments and lines (Figure S4, Supplementary Calculations). Analysis of two wild-type data sets revealed negligible and non-significant negative slopes, whereas all the transgenic lines showed negative slopes that were significant except for line 29 (Figure 3E, Figure S4C). Notably, the ME_BS&M_ x B6 lines did not show a greater loss of label from C_4_ acids than the parental B6 line whereas both of the ME_BS_ x 29 lines showed somewhat steeper and significant negative slopes than parental line 29. Collectively these results demonstrate that decarboxylation is occurring in all of the transgenic rice lines and that the biggest enhancement over decarboxylation levels seen in the parental lines is observed in the lines that accumulate high levels of NADP-ME specifically in the BS cells.

### C_4_-acid decarboxylation, ^13^C refixation by Rubisco and flux at PPDK are detected in transgenic rice lines by malate, aspartate and pyruvate feeding

Although the pulse-chase experiments established *in vivo* evidence of both the initial carboxylation by PEPC and the decarboxylation reactions in transgenic rice lines, the decarboxylation rates estimated from the chase are minimum rates because after a short pulse the malate pool used by NADP-ME will only be partly labelled. The flux measured as ^13^C released thus underestimates absolute flux and this underestimate is larger in wild-type than in the transgenic lines because carboxylation is much lower in wild-type. To eliminate this problem, we considered alternative approaches to measure flux through the C_4_ cycle. Previous modelling ^21^ had shown that operation of a C_4_ cycle between the BS and immediately adjacent M cells would lead to small changes on leaf-level CCM metrics (e.g., compensation point, ^13^C discrimination) when this local cycle operates with fluxes like those in maize. However, we realised that the effects would be undetectable with the low *in vitro* activity of PEPC (see above, Table S1) and the low observed *in vivo* flux (Figure 3) in the current transgenic rice lines. As such, we did not quantify leaf-scale CCM performance but instead focused on developing highly sensitive, step-specific assays capable of resolving low-level flux in the predominantly C_3_ anatomical background.

We developed a complementary approach to detect decarboxylation that involved feeding ^13^C-labelled malate or aspartate to detached leaves via the transpiration stream and looking for label movement into 3PGA and other CBC intermediates (Figure 4A). As an added gain, this approach monitors CO_2_ reassimilation by Rubisco. The adopted feeding approach assumes that the ^13^C detected in 3PGA and other intermediates is moving via decarboxylation and reassimilation of ^13^CO_2_ by Rubisco, and as such it was important to design a protocol that minimizes ^13^C movement by other routes. Decarboxylation releases CO_2_ from the C4 position of malate and aspartate (Figure 4A), whereas metabolism by other pathways such as the tricarboxylic acid cycle (TCA) cycle releases CO_2_ from all four C positions. To increase selectivity for ^13^CO_2_ release by decarboxylation, we therefore supplied specifically-labelled [4-^13^C]malate rather than the commercially available [U-^13^C]malate (see Methods for details of synthesis). We also supplied commercially available [4-^13^C]aspartate. The specificity of the approach also depended on ^13^C not moving from malate into CBC intermediates via shuttling of ^13^C-labelled C-skeletons. To control for this, we fed malate or aspartate at ambient (420 ppm) CO_2_ and at 50 ppm CO_2_, which is close to the CO_2_ compensation point for C_3_ species but still supports considerable flux through PEPC. The underlying reasoning was that any ^13^C moving via decarboxylation and reassimilation into CBC intermediates would be diluted by unlabelled CO_2_ that enters the leaf and is directly assimilated by Rubisco (Figure 4A). At the compensation point, however, influx of unlabelled CO_2_ into the CBC is substantially reduced. As such, if ^13^C is moving from [4-^13^C]malate or [4-^13^C]aspartate via ^13^CO_2_ into the CBC, ^13^C enrichment in CBC intermediates should be higher in 50 ppm CO_2_ than ambient CO_2_. As a further precaution, we supplied labelled malate or aspartate to leaves for relatively short times (5, 10 and 20 min) to minimize the risk of metabolism by other pathways. We also monitored labelling of citrate to check the extent to which [4-^13^C] malate was metabolised in the TCA cycle. The supplied malate and aspartate (9 mM) had only marginal effects on the malate and aspartate content of the leaves (Supplemental calculations)

**Figure 4.**
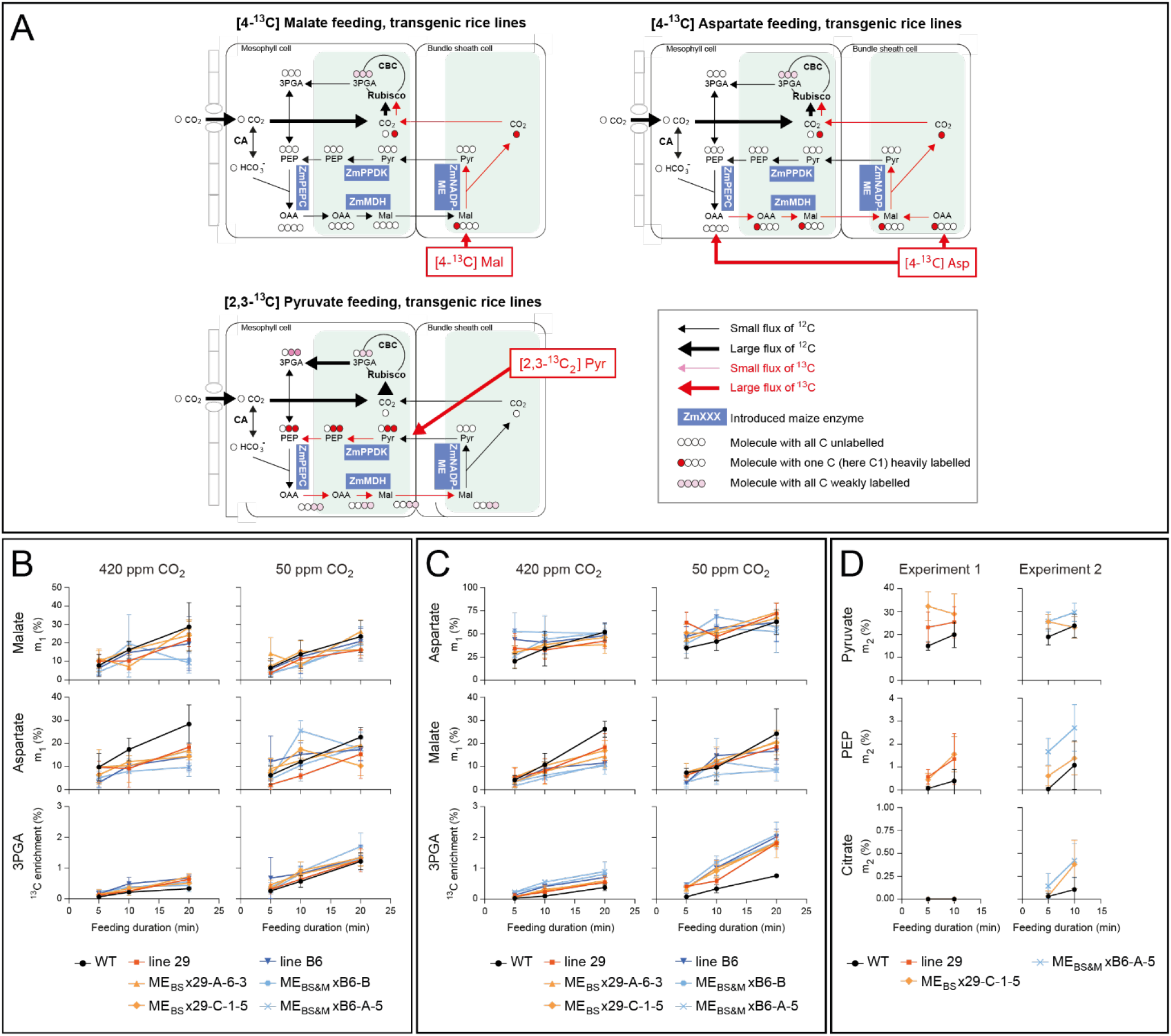
Flow of label in transgenic rice lines after malate, aspartate and pyruvate feeding experiments. **A)** Schematics to show flux after [4-^13^C]malate, [4-^13^C]aspartate or [2,3-^13^C_2_]pyruvate-feeding to transgenic rice lines. Decarboxylation and reassimilation of ^13^CO_2_ is measured as appearance of ^13^C in 3PGA and other CBC intermediates after feeding [4-^13^C]malate or [4-^13^C]aspartate. PPDK flux is measured as formation of double-labelled (m_2_ isotopologue of) PEP after feeding 2,3-^13^C_2_]pyruvate. **B)** [4-^13^C]malate-feeding and **C)** [4-^13^C]aspartate feeding experiments. In the malate- and aspartate-feeding experiments, malate and aspartate are interconverted by malate dehydrogenase and aspartate aminotransferase. The plots show fractional abundance of m*_1_* malate and m*_1_* aspartate (which provide proxies for the availability of introduced label for decarboxylation) and enrichment in 3PGA (which represents the flow of label via C_4_-acid decarboxylation and reassimilation by Rubisco). Enrichment of 3PGA depends on the flow of ^13^C via C_4_-acid decarboxylation and reassimilation, and the extent to which it is diluted by unlabelled CO_2_ entering the leaf and being fixed by Rubisco. In parallel with feeding at ambient CO_2_, feeding was performed at low-CO_2_ to test whether enrichment in 3PGA is increased when less unlabelled CO_2_ is being fixed by Rubisco. Values are mean ± SD (n = 6 for wild-type and 3 for transgenic lines). Plots showing all isotopologues of these and related metabolites are shown in Figure S5 (malate feeding) and Figure S6 (aspartate feeding). **D)** [2,3-^13^C_2_]pyruvate-feeding. The plots show the fractional abundance of m*_2_* pyruvate (which represents the availability of introduced label for PPDK) and of m*_2_* PEP (which represents the product of the PPDK reaction). The fractional abundance of m*_2_* citrate is shown as a control to indicate the extent of [2,3-^13^C_2_]pyruvate metabolism in respiratory pathways. Note that the y-axis scale is expanded compared to the other two plots. Values are mean ± SD (n = 3 to 4). Plots showing all isotopologues of these and further metabolites plus statistical analyses are shown in Figure S7 and Table S2. For original data see Supplementary Datasets 1D-I. Supported by Figures S5-S7, Table S2 and Supplementary Datasets 1D-I.

The data derived from feeding [4-^13^C]malate or [4-^13^C]aspartate to the parental B6/29 and the high ME ME_BS&M_ x B6/ME_BS_ x 29 lines are shown in Figure 4B and 4C. The fractional abundance of the m*_1_* isotopologues of malate and aspartate are shown as proxies for the extent to which labelled [4-^13^C]malate or [4-^13^C]aspartate entered the leaf. Crucially when [4-^13^C]malate or [4-^13^C]aspartate were supplied at 420 ppm or 50 ppm CO_2_, the fractional abundance of the m*_1_* isotopologues of malate and aspartate rose with time in all genotypes and conditions (Figure 4B, 4C) whereas there was almost no formation of higher isotopologues of malate or aspartate (Figure S5A, Figure S6A, Supplementary Datasets S1D-G). With the exception of m*_1_* aspartate in the [4-^13^C]aspartate-feeding experiment, which in many transgenic lines at early times showed much higher abundance than in wild-type and did not rise with time (Figure 4C), there was a trend to lower labelling of m*_1_* malate and m*_1_*aspartate in the transgenic lines than wild-type, consistent with faster decarboxylation. Secondly, the enrichment in 3PGA is shown as a proxy for the rate of ^13^CO_2_ reassimilation by Rubisco relative to the influx of unlabelled CO_2_ (see Supplementary Calculations). Notably 3PGA enrichment rose more rapidly in the transgenic lines than in wild-type (Figure 4B, 4C). Further evidence for increased refixation of ^13^CO_2_ was provided by the observation that other CBC intermediates and related intermediates had higher enrichment in the transgenic lines than wild-type (Figure S5B, S6B), as expected because label fixed into 3PGA moves rapidly into other CBC intermediates ^20,22,23^. Crucially, enrichment in 3PGA and other CBC intermediates rose more quickly when feeding was performed at 50 ppm CO_2_ as opposed to feeding at 420 ppm CO_2_. Finally, labelling in other organic acids including fumarate, citrate and pyruvate was determined to assess how much metabolism of [4-^13^C]malate and [4-^13^C]aspartate was occurring by routes other than decarboxylation (Figures S5B, S5C, S6B, S6C; Supplementary Datasets S1D-G). There was negligible labelling of citrate, indicating that [4-^13^C]malate was not entering the TCA cycle via mitochondrial pyruvate dehydrogenase (PDH). Enrichment in pyruvate was low and showed no consistent differences between wild-type and the transgenic lines. Note that because there was often substantial variation between replicates in these feeding experiments, we developed a method to reduce this variability based on co-variation between labelling of different metabolites (see Table S2 for details). In combination, the results obtained demonstrate, firstly, that C_4_ acid decarboxylation is significantly faster in the transgenic lines than in wild-type rice (although rates did not appear to differ in parental (low ME) versus ME_BS&M_ x B6 and ME_BS_ x 29 (high ME) lines) and, secondly, that released ^13^CO_2_ is being reassimilated by Rubisco.

To determine whether the final step of the C_4_ pathway, was functional in transgenic rice lines, flux at PPDK was investigated by feeding labelled pyruvate and monitoring conversion to PEP. This approach depended on minimizing movement of label from pyruvate to PEP via PDH because any ^13^CO_2_ released by PDH from pyruvate could be reassimilated by Rubisco to produce labelled 3PGA, from which label could move into PEP. To minimize release of ^13^CO_2_ by either mitochondrial or plastidial PDH, we supplied leaves with commercially-available [2,3-^13^C_2_]pyruvate (Figure 4A). We also focused our analysis on fractional abundance of the m*_2_* isotopologue of PEP, the form that would be produced by PPDK from [2,3-^13^C_2_]pyruvate. A lower concentration (2mM) was supplied than for malate or aspartate to avoid any large elevation of leaf pyruvate content (Supplemental calculations). Two experiments were performed using combinations of wild-type rice, parental line 29, ME_BS_ x 29-C-1-5 and ME_BS&M_ x B6-A-5. In both experiments, the fractional abundance of the m*_2_* isotopologue of pyruvate rose to 20-30% (Figure 4D) and as expected accounted for the vast majority (>95%) of the labelled pyruvate (Figure S7A, Supplementary Datasets S1H-I). There was a trend to a higher fractional abundance of m*_2_* pyruvate in the transgenic lines, but this was not significant except for line ME_BS_ x 29-C-1-5 in experiment 1 (Figure S7B). The fractional abundance of the PEP m*_2_*isotopologue was higher in the transgenic lines than wild-type rice (Figure 4D, Figure S7B), and as expected if PPDK is converting m*_2_* pyruvate into m*_2_* PEP, the fractional abundance of m*_2_* PEP was higher than that of the m*_2_* isotopologue of 3PGA or other CBC intermediates (Figure S7A-B). The lower abundance of m*_1_* PEP than m*_2_* PEP or m*_1_* 3PGA (Figure S7A-B) is consistent with the m*_1_* isotopologues being formed at a low basal rate, independently of the introduced ZmPPDK. A plausible route would be via release of ^13^CO_2_ and refixation by Rubisco into 3PGA. Collectively, therefore, these observations are consistent with slow movement of ^13^C from [2,3-^13^C]pyruvate to 3PGA via routes other than PPDK in all of the genotypes, but also with an enhancement of flux via PPDK in some of the transgenic lines.

### Quantitative estimation of fluxes

Although the data presented so far provide qualitative evidence for enhanced fluxes at the introduced enzymes in the transgenic lines, information about absolute flux is required to compare *in-vivo* flux with the introduced enzymatic capacity (as measured in an *in-vitro* assay) and with the fluxes that would be required for C_4_ photosynthesis. Flux estimates are provided in Figure 5 (for details of how flux was estimated see legend of Figure 5, Supplementary Table S2 and Supplementary Calculations). Notably the estimates of flux at PEPC will be close to actual *in vivo* flux. For NADP-ME, estimates of flux from pulse chase experiments will be underestimates because the malate pool is only partly labelled after the short 15-30 s pulse, with underestimation especially large for wild-type. Estimates of NADP-ME activity (and CO_2_ re-assimilation by Rubisco) will be more realistic when based on enrichment in 3PGA in [4-^13^C]malate-feeding experiment and [4-^13^C]aspartate-feeding experiments. However, they may be overestimates or underestimates, depending on whether the malate pool used by NADP-ME is more or less heavily-labelled than the overall pool of malate or aspartate in the leaf. Flux at PPDK may be underestimated due to restricted access of supplied [2,3-^13^C_2_]pyruvate to PPDK in the chloroplasts of M cells.

**Figure 5.**
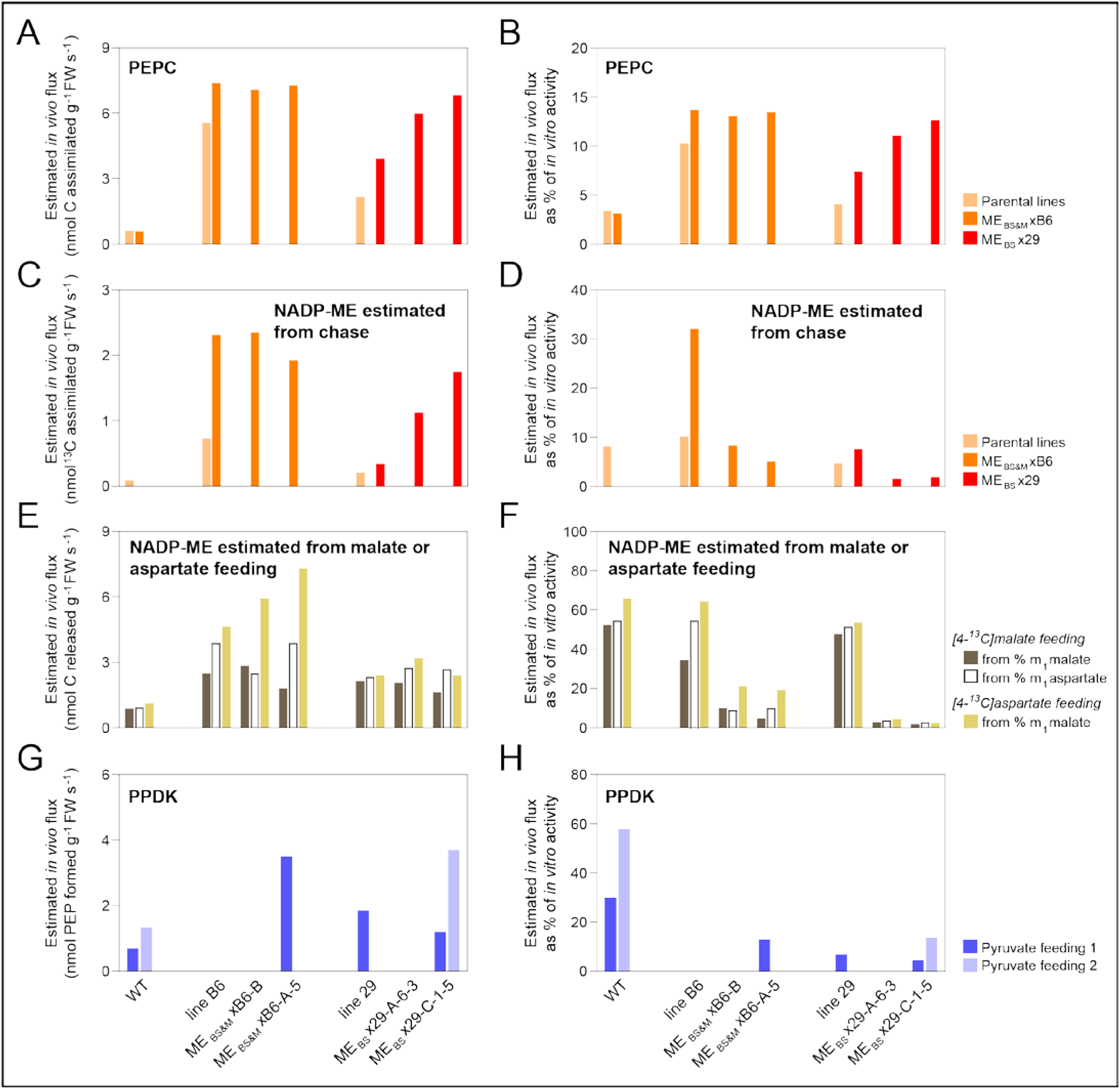
Summary of estimated *in vivo* fluxes at PEPC, NADP-ME and PPPK in transgenic rice lines. **A)** *In vivo* flux at PEPC estimated from ^13^CO_2_ pulse experiments as the sum of the abundance of labelled malate and aspartate isotopologues (Figure 3B), with a small upwards correction for ^13^CO_2_ released by decarboxylation of malate and aspartate during the pulse. Given that there is little dilution of the incoming ^13^CO_2_, estimated ^13^C fixation approximates to absolute flux, nmol C_4_ acid g^-1^ FW s^-1^. **B)** *In vivo* flux as a percentage of PEPC activity measured *in vitro*, i.e., enzymatic capacity (taken from Supplementary Table S1). **C)** *In vivo* flux at NADP-ME estimated from pulse-chase experiments, using the slope of the plot of summed corrected m*_1_* malate and aspartate *versus* time (Figure 3D-E, Figure S4), with a small upward correction to include ^13^C release from the m*_2_*, m*_3_* and m*_4_* isotopologue (see Supplementary Calculations). The estimate has the unit nmol ^13^C released g^-1^ FW s^-1^ and underestimates absolute flux because the malate pool is only partly labelled after the short 15-30 s pulse. **D)** *In vivo* flux at NADP-ME estimated from the pulse-chase experiment as a percentage of NADP-ME activity measured *in vitro*. **E)** Estimation of *in vivo* flux at NADP-ME was from enrichment in 3PGA in the [4-^13^C]malate-feeding and [4-^13^C]aspartate-feeding experiments.^13^C enrichment in 3PGA will depend i) on the rate of ^13^CO_2_ release by NADP-ME and ^13^CO_2_ refixation by Rubisco, and ii) the extent to which this ^13^CO_2_ is diluted by ^12^CO_2_ that is fixed by Rubisco. Slopes of plots of 3PGA enrichment versus fractional abundance of the m*_1_* isotopologue of malate reflect or aspartate are summarized in Table S2, and plots are provided in Supplementary Calculations. Net photosynthesis by detached wild-type rice leaves in the conditions used in the feeding experiments was about 76 nmol CO_2_ g^-1^FW s^-1^. Photosynthesis rates of several of the transgenic lines are similar to wild-type rice ^18,19^. Assuming that a 4:1 ratio of carboxylation to oxygenation would be equivalent to gross flux of about 86 nmol CO_2_ g^-1^FW s^-1^ at Rubisco, multiplying the slope of a plot of 3PGA enrichment *versus* fractional abundance of the m*_1_* isotopologue of a C_4_ acid (Table S2) by 86 nmol CO_2_ g^-1^FW s^-1^ gives an estimate of the rate of decarboxylation (nmol malate g^-1^ FW s^-1^). The panel shows three flux estimates. Two estimates were made from the [4-^13^C]malate-feeding experiment, using the m*_1_* fractional abundance of malate or the m*_1_* fractional abundance of aspartate as alternative proxies for the m*_1_* fractional abundance in the malate pool that is the substrate for NADP-ME. A third estimate was made from the [4-^13^C]aspartate-feeding experiment, using the m*_1_*fractional abundance of malate as a proxy for the m*_1_* fractional abundance in the malate pool that is the substrate for NADP-ME. The approach assumes that the overall m*_1_* fractional abundance of a given isotopologue is a proxy for the pool of malate that is the substrate for NADP-ME, and none of the proxies are perfect. Estimates from the [4-^13^C]aspartate-feeding experiment based on m*_1_*aspartate were excluded; its unusual labelling kinetics (Figure 4C) resulted in low flux estimates that were considered to be outliers. Note the y-axis scale has a 3-fold greater range than panel C. **F)** *In vivo* flux at NADP-ME estimated from [4-^13^C]malate-feeding and [4-^13^C]aspartate-feeding as a percentage of NADP-ME activity measured *in vitro*. **G)** Flux at PPDK estimated from [2,3-^13^C_2_]pyruvate-feeding experiments.. To estimate flux (nmol pyruvate g^-1^ FW s^-1^), the slope of the plot of the fractional abundance of the m*_2_* isotopologue of PEP versus the fractional abundance of the m*_2_* isotopologue of pyruvate (Table S2) was multiplied by 86 nmol g^-1^FW s^-1^, and in this case divided by two because each molecule of [2,3-^13^C_2_]pyruvate contains two atoms of ^13^C. The conversion to absolute flux assumes that the overall m*_2_* fractional abundance of pyruvate reflects that in the pool that is the substrate for PPDK. **H)** *In vivo* flux estimated from 2,3-^13^C_2_]pyruvate-feeding as percentage. of PPDK activity measured in vitro. For details see Supplementary Calculations. Supported by Table S2 and Supplementary Calculations.

*In vivo* flux at PEPC was estimated from ^13^CO_2_ pulse experiments as the sum of the abundance of labelled malate and aspartate isotopologues, with a small upwards correction for ^13^CO_2_ released by decarboxylation of malate and aspartate during the pulse. Values in different lines and experiments ranged from 2.2-7.4 nmol g^-1^ FW s^-1^ (Figure 5A) and were higher in the ME_BS&M_ x B6 lines than in the ME_BS_ x 29 lines. In all cases, *in vivo* flux was much lower than PEPC activity in *in vitro* assays (4.1-13.6% of introduced enzyme capacity) (Figure 5B). *In vivo* flux at NADP-ME estimated from the pulse-chase labelling experiment (Figure 5C) ranged from 0.21-2.3 nmol ^13^C g^-1^ FW s^-1^. There was a trend to higher detected flux in the ME_BS_ x 29 lines than parental line 29 but flux was similar in the ME_BS&M_ x B6 and parental B6 lines. Except for one experiment with line B6, the estimated fluxes were <10% of the introduced NADP-ME capacity, and were especially low (<2%) in the high ME_BS_ x 29 lines (Figure 5D). NADP-ME flux estimates from the [4-^13^C]malate-feeding experiment, using abundance of m_1_ malate and m_1_ aspartate as alternative proxies for labelling of the malate pool that serves as a substrate for NADP-ME, and from the [4-^13^C]aspartate-feeding experiment using abundance of m_1_ malate (Figure 5E) ranged between 1.8-7.3 nmol g^-1^ FW s^-1^. These are also estimates for the rate of CO_2_ reassimilation by Rubisco. Note that there was no consistent difference between the parental lines and ME_BS_ x 29 and ME_BS&M_ x B6 lines. Note also that estimates using m*_1_*aspartate were considered unreliable due to the unusual labelling kinetics of m*_1_* aspartate when [4-^13^C]aspartate was fed (Figure 4C; for further reasons see Supplementary Calculations). In all cases, the estimated *in vivo* flux at NADP-ME was lower than NADP-ME *in vitro* activity; moderately in the parental B6 and 29 lines (35-64% of introduced enzyme capacity) but strongly in the ME_BS&M_ x B6 (5-21%) and ME_BS_ x 29 (1.8-4.4%) lines (Figure 5F). This indicates that the 6-fold increase in introduced NADP-ME capacity in the ME_BS&M_ x B6 lines and the 30-fold increase in the ME_BS_ x 29 lines (Table S1) was not being converted into meaningful increases of estimated *in vivo* flux. Finally, flux at PPDK was estimated from [2,3-^13^C_2_]pyruvate-feeding experiments (see details in legend of Figure 5 and Supplementary Calculations). In this case, estimated flux ranged from 1.2 -3.7 nmol g^-1^ FW s^-1^ (Figure 5G) and was again lower (4.4-14%) than the introduced enzyme capacity (Figure 5H).

Collectively, the results demonstrate *in vivo* flux at all key reactions of the C_4_ cycle in an engineered C_3_ plant.

## DISCUSSION

### C_4_ fluxes detected after the implementation of new sensitive methods

The ability to measure *in-vivo* flux is a prerequisite for any attempt to engineer a metabolic pathway. We have demonstrated that the expression of maize genes encoding C_4_ enzymes in rice results in substantial flux enhancement at all four core steps of the C_4_ cycle, consistent with activity of the introduced enzymes. The estimated *in vivo* fluxes are often only a small fraction of the introduced enzymatic capacity (Figure 5B, 5D, 5F, 5H), but the estimated *in vivo* fluxes are approximate and probably minimum estimates.

Previous attempts to engineer C_4_ cycle enzymes into rice and other C_3_ species succeeded in introducing the proteins but had little or no impact on detectable flux *in vivo*. This was partly because the introduced activities were low, but was mainly complicated by difficulties in detecting low-level flux against a background of rapid C_3_ photosynthesis, and by the anatomical background in which the introduced enzymes operate. Figure 6 (adapted from ^21^) illustrates this complication by showing the C_4_ biochemical pathway introduced to rice and the localization of the maize enzymes in this anatomical background. Because rice has 6 to 9 mesophyll cells between vascular bundles whereas maize and sorghum typically have 2, there are fewer BS cells in rice on a leaf area or fresh weight basis. As such, activities of enzymes accumulating specifically in the rice BS will appear to be lower on a whole leaf basis than in C_4_ leaves, even if the activity per BS cell is similar. Similarly, measurements of flux at enzymes accumulating in M cells will often only represent activities in M cells adjacent to the BS.

**Figure 6.**
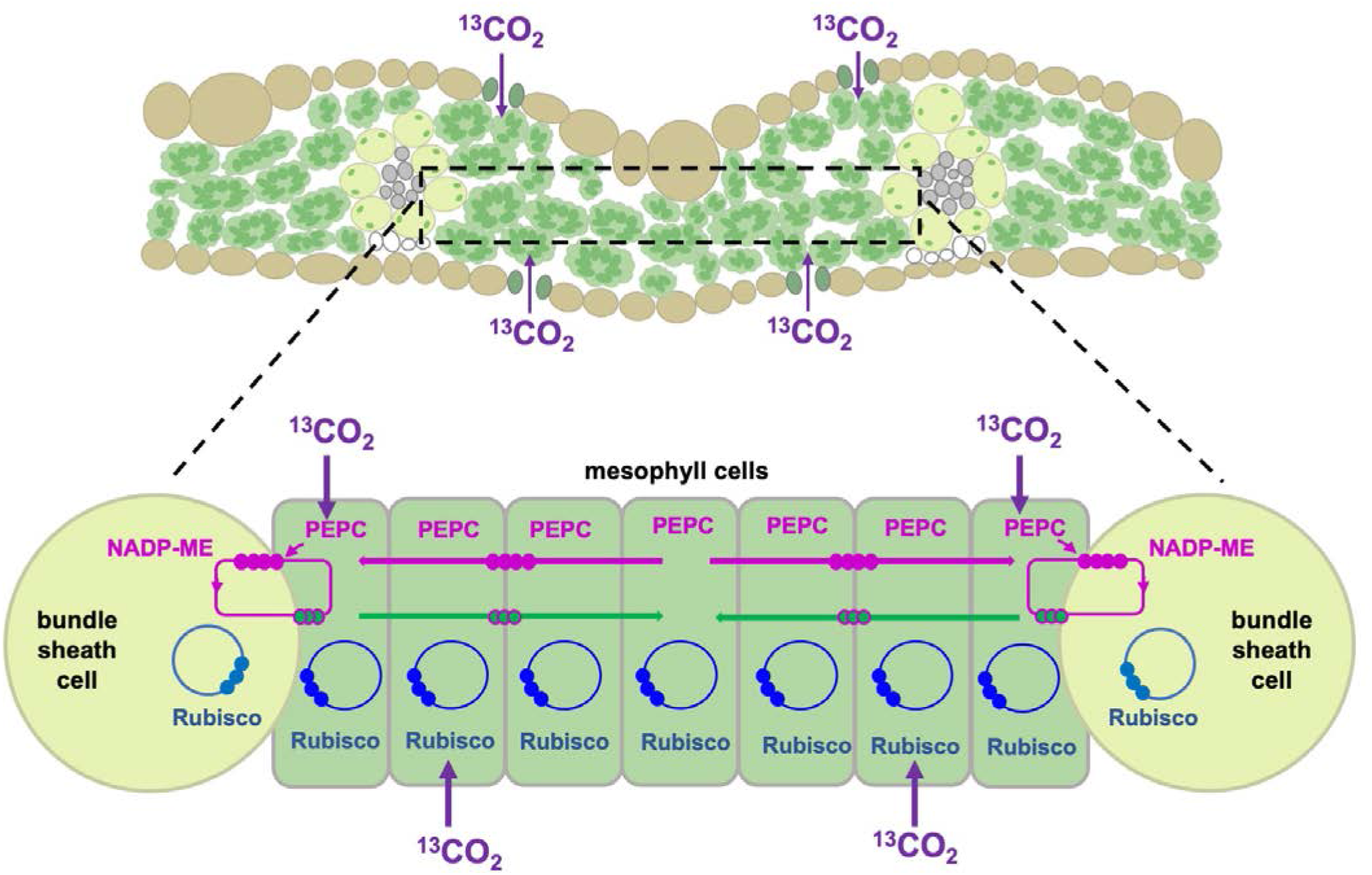
Schematic of C_4_ pathway introduced to rice. **A)** Transverse sections of C_3_ leaves reveal veins that are encircled by BS (light green) that are typically separated by 6 to 9 M cells (dark green). Unlike C_4_ leaves, chloroplasts are much more abundant in M cells than BS cells. **B)** In the ME_BS_ x 29 lines, PEPC is expressed in the M cells between the vascular bundles, co-located with CA, NADP-MDH, PPDK and the endogenous rice Rubisco plus the CBC (blue pathway). In these plants, against a large background of CO_2_ fixation by Rubisco, PEPC fixes bicarbonate to produce C_4_ acids which must then diffuse to the BS cells to be decarboxylated (magenta pathway). Pyruvate (green arrows) must diffuse back to the M cells to regenerate PEP to supply PEPC. The diagram was redrawn from ^21^, where implications for quantitative fluxes were also discussed. It is proposed that the majority of the C_4_ pathway activity in these plants occurs around the veins where rapid intercellular diffusion of metabolites is more likely ^21^.

Consideration of the factors discussed above enabled us to develop methods that were sensitive enough to detect C_4_ fluxes against a background of C_3_ anatomy and photosynthesis. The methods allowed direct detection of tangible *in-vivo* fluxes at four key steps around the C_4_ cycle: initial C fixation into C_4_ acids by PEPC; C_4_-acid decarboxylation via NADP-ME; reassimilation of released ^13^CO_2_ by Rubisco; and PEP regeneration by PPDK. Although not explicitly tested, the rapid labelling of both malate and aspartate in the pulse and interconversion of malate and aspartate in feeding experiments provided indirect evidence for *in-vivo* activity of ZmNADP-MDH. We also developed straightforward approaches to estimate absolute flux, based on the labelling kinetics of the substrates and immediate products of the reactions (Supplementary Calculations). These calculations are particularly suitable for analysis of non-steady state labelling kinetics, which were unavoidable in the short-term labelling experiments that were essential to detect fluxes in the C_4_ cycle. The method also allows problems due to metabolite compartmentation to be explicitly addressed and the likely resulting error assessed.

Despite their sensitivity and utility, the protocols we have developed have limitations. Metabolite compartmentation in particular, especially of malate ^20,24-26^, poses a major complication. This does not seriously affect estimation of flux at PEPC, which is cytosolic, but does impinge on estimation of C_4_-acid decarboxylation by chloroplastic NADP-ME, and potentially also PPDK. That said, despite some uncertainties, similar estimates of the rate of C_4_-acid decarboxylation in the transgenic lines were provided by two completely independent types of experiment and calculation - pulse-chase experiments and [4-^13^C]malate feeding. A potential issue with the feeding approach is that the rate of supply of specifically-labelled [4-^13^C]malate or [4-^13^C]aspartate to the BS cells may limit the rate of decarboxylation. This is particularly pertinent because the xylem parenchyma symplast is poorly connected to the phloem and surrounding vasculature in rice ^27^. Limited supply of labelled malate and aspartate could thus limit the maximum detectable flux through NADP-ME, especially in the ME_BS_ x 29 lines. Going forward, limitations to uptake rates of malate and aspartate in feeding experiments may become more apparent in lines with higher C_4_ cycle fluxes, and thus modifications in the protocol to increase uptake may be required. Alternatively, once higher fluxes at PEPC are achieved, analysis of pulse-chase experiments may suffice to monitor *in vivo* flux at NADP-ME.

### Limitations to C_4_ photosynthetic flux in transgenic rice lines

The improved labelling methodology reported above clearly indicates low but demonstrable flux through all C_4_ pathway steps in the transgenic rice lines produced. Estimated *in vivo* fluxes for NADP-ME and PPDK in transgenic lines (Figure 5E, 5G) exceed *in vitro* levels of activity detected in wild-type samples (Table S1), consistent with the elevated fluxes being due to the introduced enzyme. Although the same is not true for PEPC, several factors restrict operation of maize PEPC in rice (see below).

The estimated in vivo fluxes were low compared to those required for an effective CCM. In a recent ^13^CO_2_ pulse-chase study with four C_4_ species including maize ^28^, after a 10 s pulse the m*_1_* isotopologues of malate and aspartate accounted for >50% of the photosynthetic pool of malate and aspartate, and about half were lost within ∼10 s in the chase, with estimated rates of carboxylation and decarboxylation in the range of 103-179, and 65-150 nmol ^13^C g^-1^ FW s^-1^, respectively. This contrasts with the transgenic rice lines where, after a 60 s pulse, the m*_1_* isotopologues account for only ∼10% of the malate and aspartate pool (Fig. 3B), and in the chase (about half are lost after 50-80 s (after correction for continuing labelling for PEP, Figure 3D, Supplementary Figure S3B), with estimated rates of carboxylation and decarboxylation of the order of 4-6 and 1-2 nmol ^13^C g^-1^ FW s^-1^ (Figure 5), Overall, the estimated in vivo fluxes in the rice lines at PEPC, NADP-ME and PPDK (Figure 5) lie in the range of 2-7%, 4-10% and 1-3% of those required in C_4_ plants ^28^. They were 3-9, 4-9 and 2-5% of the ambient rate of CO_2_ fixation in rice (∼76 µmol m^-2^ s^-1^).

Further improvements in flux in rice will required higher expression of PPDK and, especially, PEPC. Use of a genomic PEPC sequence enhanced PEPC expression ^29^ compared to the cDNA sequence used here and in ^21^ but by only ∼50% and further large gains in expression are still required. It will also require better utilisation of the introduced enzyme capacity. For all enzymes, the estimated *in vivo* flux in the transgenic rice lines, even when catalysed by feeding labelled photosynthetic intermediate, was considerably lower than introduced capacity (Figure 5B, 5D, 5F, 5H). It has been demonstrated that several factors restrict in vivo activity of maize PEPC in rice ^29^, including known phosphorylation mechanisms ^30,31^. *In vivo* activity of NADP-MDH may be similarly restricted by lack of thioredoxin-mediated reduction ^32,33^, and of PPDK by lack of phosphorylation/dephosphorylation ^34-37^. *In vivo* fluxes may also be restricted by low endogenous activities of metabolite transporters. Although the transporters required for the C_4_ cycle are present in C_3_ plants, transcriptome analyses indicate that they are upregulated 10-100-fold in C_4_ plants ^38^. Malate transport into BS chloroplasts by endogenous rice transporters may restrict flux at NADP-ME both in pulse-chase experiments and when ^13^C malate is fed to leaves. Similarly, restricted pyruvate export from BS chloroplasts and pyruvate and PEP transport in M chloroplasts likely constrains PPDK, NADP-MDH and PEPC activities. There are thus a number of additional metabolic modifications that could be made to improve *in vivo* flux levels.

The capacity for C_4_ cycle function in rice will be inherently constrained by C_3_ leaf anatomy and in addition, the anatomical differences between rice and C_4_ plants will lead to *in vivo* flux being underestimated in the transgenic lines. For example, sorghum has 10 veins per linear mm across the mature leaf blade on average whereas rice has only 5.5 ^39^, giving a “correction factor” of 1.8. The factor will be species dependent and also depend on cell size but because of it, *in vivo* fluxes at NADP-ME are certainly underestimated in rice relative to those in maize or sorghum. For PEPC and PPDK, a numerical correction factor is not as important, given the large number of mesophyll cells in any given area of the rice leaf blade. However, the diffusion limitations imposed by vein spacing may still be relevant. During ^13^CO_2_ pulse chase experiments, malate or aspartate produced by PEPC in cells at a distance from the veins must diffuse back to the BS for decarboxylation, at least in the ME_BS_ x 29 lines. In C_4_ plants, plasmodesmata (PD) connections between the M and BS cells are greatly enhanced to maximise this diffusive process but in rice, PD density between M cells is an order of magnitude lower ^40^. It is thus likely that the majority of C_4_ flux detected in ^13^CO_2_ pulse chase experiments is occurring in M cells adjacent to the BS (Figure 6). Even in ^13^C feeding experiments, detection of flux at PPDK will depend on how efficiently the fed pyruvate gets into the M cells (presumably either through the cell wall in the transpiration stream and then across the M plasma membrane, or into the symplasm near the xylem and then from cell to cell by PD). A true picture of the capacity for *in vivo* flux at C_4_ enzymes in rice will therefore only be obtained when the enzymes are introduced into a modified anatomical chassis.

### Prospects for C_4_ rice

Now that we have established that the four main reactions needed for the biochemical CO_2_ pump operate in rice, they will need to be optimised. Other traits relevant to efficient operation of the C_4_ cycle will also need to be modified. Although extractable activities of NADP-ME and NADP-MDH have been achieved that are in the same order as those from C_4_ plants, PEPC and PPDK activities have only reached 5% and 15% of maize levels respectively. A variety of transcriptional and translational enhancement strategies are underway to increase enzyme activity and it may also be necessary to adjust post-translational regulation (see above). The ^13^C flux-detection protocols described here will allow direct testing of the efficacy of these strategies and will also be invaluable in determining *in vivo* flux in lines with altered leaf anatomy; it is likely that both leaf vein density and plasmodesmatal frequencies between M and BS cells will need to be increased.

## MATERIALS and METHODS

### Rice germplasm

Experiments were performed using wild-type rice (*Oryza sativa spp. japonica*) cv. Kitaake and transgenic lines generated in this background. Parental lines B6 and 29, carrying a multigenic construct for expression of maize carbonic anhydrase, PEPC, NADP-MDH, NADP-ME and PPDK have been described previously ^18^.

### Plant growth conditions

For genotyping, molecular characterization and propagation, rice plants were grown in a controlled environment chamber (Model PGC Flex, Conviron, Winnipeg, MB, Canada) with a 16h/8h photoperiod and an irradiance of 400 μmol m^-2^ s^-1^ supplied by a mixture of fluorescent tubes (Master TL5 HO 54W/840, Philips Lighting, The Netherlands) and halogen incandescent globes (42 W 2800 K warm clear glass 630 lumens, CLA, Brookvale, Australia). The day/night temperatures were 28°C/22°C and the relative humidity was a constant 60%. Plants were individually grown in 1L pots in a soil mix composed of 80% peat, 10% perlite and 10% vermiculite (pH 5.6-5.8) mixed with 5 g slow-release fertilizer (Osmocote, Evergreen Garden Care, Australia) supplied at the beginning of the growth cycle. All pots were kept at field water capacity.

For ^13^CO_2_ labelling and ^13^C-feeding experiments, seeds were germinated on agar and then transplanted to 1L pots filled with a 2:1 mixture of peat and medium-sized grain quartz sand (Einheitserdewerke Werkverband e.V, Sinntal-Altengronau, Germany), supplemented with 0.66 mL L^-1^ Plantacote Depote 4M (Wilhelm Haug GmbH & Co. KG, Ammerbuch, Germany) and 0.66 mL L^-1^ Fetrilon Combi (COMPO EXPERT GmbH, Muenster, Germany) slow-release fertilizer granules. The pots were submerged in water. Plants were grown in a controlled environment chamber with a 14h/10h photoperiod and an irradiance of 560 µmol m^-2^ s^-1^ supplied by white and far-red LEDs tuned to approximate a sunlight-like spectrum. The day/night temperatures were 28°C/20°C and relative humidity was a constant 80%. Rice plants were used for ^13^C-labelling experiments at 4-6 weeks after germination.

### Introgression of a maize NADP-ME gene construct into lines B6 and 29

Transgenic lines carrying multigenic constructs for BS cell-specific expression of the maize NADP-ME and a β-glucuronidase (GUS) reporter were generated as described in ^19^ (Figure S1A). One construct comprised: (i) a synthetic gene encoding a designer transcription activator-like effector (dTALE1), under the control of the *Zoysia japonica PEP CARBOXYKINASE* promoter; (ii) a *ZmNADP-ME* gene construct (including synthetic introns, 5′ KOZAK sequence and 3′-UTR) under the control of a dTALE1-cognate synthetic TALE-activated promoter (STAP 1.45); and (iii) a *GUS* gene construct under the control of a dTALE1-cognate STAP1.1 promoter (Figure S1A). The second construct comprised: (i) a synthetic gene encoding a designer transcription activator-like effector (dTALE1), under the control of the *Zoysia japonica PEP CARBOXYKINASE* promoter; (ii) a *ZmNADP-ME* gene construct under the control of a dTALE1-cognate synthetic TALE-activated promoter (STAP 1.3); (iii) a *GUS* gene construct under the control of a dTALE1-cognate STAP1.1 promoter; (iv) a NLS-eGFP gene construct under the control of a dTALE1-cognate STAP1.4 promoter; and (v) a NLS-kOrange gene construct under the control of a dTALE1-cognate STAP1.5 promoter (Figure S1A).

Two independent dTALE-STAP ME lines were established for crossing with lines B6 and 29 that carry a multigenic construct encoding a minimal set of five C_4_ pathway enzymes ^18^. Line 29 (♂) was crossed with dTALE-STAP T1 line ME 3 (♀) which had a single insertion of construct 1 and line B6 (♂) was crossed with dTALE-STAP T1 line 1 (♀) which had multiple-insertions of construct 2. Crossing was performed by manual emasculation of the developing florets from the maternal parent followed by pollination from the paternal parent according to ^41^. F1 seeds were germinated in transparent magenta boxes containing solid MS medium supplemented with 50 mg L^-1^ of hygromycin for 2-3 weeks at 25°C with a 16 h light/8 h dark cycle. Seedlings were transferred to soil in the controlled environment chamber as described above. F1 plants were first genotyped to select for successful crosses and then screened for the presence of the five C_4_ genes based on transcript and protein expression. Selected F1 plants with confirmed expression were selfed. The copy number of the *hygromycin phosphotransferase* (*hpt*) selectable marker gene in F2 individuals was determined by digital droplet PCR (IDna Genetics Ltd., Norwich, UK) to identify homozygous lines.

### Genotyping and transcript detection

Leaf discs were collected from the mid-distal leaf blade portion of the youngest fully expanded leaf from the central shoot of 4-week-old rice plants, frozen in liquid N_2_ and stored at minus 80°C. Frozen samples were homogenised using a Qiagen TissueLyser II (Qiagen, Hilden, Germany; https://www.qiagen.com). RNA was extracted using an RNeasy Plant Mini Kit (Qiagen) and genomic DNA was isolated from the same sample by collecting the flow through from the spin column for subsequent PCR. PCR genotyping was performed on genomic DNA using primer pairs for detection of (i) the five-enzyme construct from the B6 or 29 parental lines: 5′-CGTCATACAAGCCGGCAATG and 5′-TGTGCTTCCAGACTCTGCAG (5s or B6) and (ii) the dTALE-STAP ME construct: 5′-GCCGTTCACCTCCATCAGAA and 5′TTGATTTCACGGGTTGGGGT.

For RT-qPCR, DNA was removed from the RNA preparation using an Ambion TURBO DNA free kit (Thermo Fisher Scientific, Tewksbury, MA; https://www.thermofisher.com), and RNA quality was determined using a NanoDrop (Thermo Fisher Scientific). One microgram of RNA was reverse transcribed into cDNA using SuperScriptTM III Reverse Transcriptase (Thermo Fisher Scientific). Real-Time qPCR was performed on the cDNA using a T100TM Thermal Cycler (Bio-Rad, Singapore; https://www.bio-rad.com/) using GoTaq® Green Master Mix (Promega, Australia; https://www.promega.com). Primer pairs were designed using Primer3 in Geneious R9.1.1 (https://www.geneious.com) and used to quantify transcripts from the five maize C_4_ pathway transgenes and the rice *ELONGATION FACTOR 1α* (*OsEF1α*) reference gene:

*ZmCA* – 5′-CATTCACCGTCCGCAACATC and 5′-AGGCGAACACCATGTACCTG

*ZmPEPC* – 5′-CGTCATACAAGCCGGCAATG and 5′-TGTGCTTCCAGACTCTGCAG

*ZmNADP-MDH* – 5′-GCCCCTCTCGGCCGC and 5′-CACCTTCGAGGGCTTGAAACG

*ZmNADP-ME* – 5′-GCCGAGCAGACCTACTTGTT and 5′-GTTGCGGTACACCGGCG

*ZmPPDK* – 5′-GAATCCCAGAGCATCCCGAG and 5′-GTGCAGGACAGGGAAACGTA

*OsEF1 α* – 5′-TCTTCCTGGTGACAACGTCG and 5′-TGGGCTTGGTGGGAATCATC

The expected sizes of the amplicons were confirmed by electrophoresis on 1% (w/v) agarose gels.

### Protein quantification by immunoblotting

Leaf discs (approx. 0.4 cm^2^) were collected from the mid-distal region of the youngest fully expanded leaves of 4-week-old plants, immediately frozen in liquid N_2_ and kept at -80°C until analysis. Leaf discs were homogenized in ice-cold extraction buffer containing: 50 mM 4-(2-hydroxyethyl)-1-piperazinepropanesulphonic acid-NaOH, pH 7.8, 5 mM MgCl_2_, 2 mM EDTA, 5 mM dithiothreitol, 1% (w/v) polyvinylpolypyrrolidone, 0.1% (v/v) Triton X-100, and 1% (w/v) protease inhibitor cocktail (Sigma-Aldrich, St Louis, MI, USA, catalogue No. P9599; https://www.sigmaaldrich.com). Extracts were supplemented with 2% (w/v) sodium dodecyl sulphate and incubated at 65 °C for 10 min. Proteins in the extract were separated by polyacrylamide gel electrophoresis (Nu-PAGE 4-12% Bis-Tris gel, Invitrogen, Life Technologies Corporation, Carlsbad, CA) and then electroblotted onto a PVDF membrane using a wet transfer method. After blocking, the membrane was probed with specific antibodies against *Zm*PEPC (1:10,000 dilution; ^16^), *Zm*PPDK (1:20,000 dilution; ^19^), *Zm*NADP-MDH (1:5000 dilution; ^19^), *Zm*ME (1:5000 dilution; ^42^), and the AcV5 tag (1:10,000 dilution; Abcam, Cambridge, UK, catalogue No. ab49581; https://www.abcam.com) and detected as in ^43^.

### Immunolocalisation of maize C_4_ enzyme proteins in rice leaves

For fluorescent immunodetection of proteins, leaf tissue was cut directly into fixing solution (4% (w/v) paraformaldehyde, 0.2% (w/v) glutaraldehyde, 0.01% (v/v) Tween-20, 25 mM sodium phosphate buffer, pH 7.2), vacuum-infiltrated until the tissue was submerged, then transferred into fresh fixative solution and incubated for 3-4 h at 4 °C. After rinsing in 25 mM sodium phosphate buffer, thin leaf sections were hand-cut using a razor blade and placed into blocking solution (20 mM 2-amino-2- (hydroxymethyl)-1,3propanediol, 154 mM NaCl, 0.1% (v/v) Tween 20, 3% (w/v) dried milk powder). Sections were incubated overnight with one of the primary antibodies (see above for the sources of the antibodies) in blocking solution: *Zm*PEPC (1:1000 dilution), *Zm*PPDK (1:100 dilution), *Zm*MDH (1:100 dilution), *Zm*ME (1:100 dilution), or AcV5 tag (1:100 dilution). For visualization, sections were incubated with AlexaFluor 488-conjugated goat anti-rabbit antibody (Thermo Fisher Scientific, catalogue No. A-11070; 1:200 dilution) or, for AcV5, with AlexaFluor 488-conjugated goat anti-mouse antibody (Abcam, catalogue No. ab150117; 1:200 dilution) for 2 h in the dark, and then treated for 5 min with 0.05% (w/v) calcofluor white to stain cell walls. Sections were examined with an LSM780 UV-NLO confocal microscope (Zeiss, Oberkochen, Germany; https://www.zeiss.com), and fluorescence signal was collected at 546–600 nm for Alexa Fluor 488 (excitation 488 nm), 434–445 nm for cell walls (excitation 405 nm) and 650–742 nm for chlorophyll autofluorescence (excitation 633 nm). Images were processed using ImageJ software (Rasband, W.S., ImageJ, U. S. National Institutes of Health, Bethesda, Maryland, USA, https://imagej.net/ij/, 1997-2018).

### Malic enzyme assay

NADP-ME activity was assayed spectrophotometrically in rice leaf extracts as described in ^44^.

### Pulse and pulse-chase labelling of rice leaves

Rice leaves were labelled with ^13^CO_2_ as described in ^18^ and in ^28^. Labelling was performed within the growth chamber (irradiance 560 µmol m^-2^ s^-1^, 28°C) between 2 to 8 hours after the start of the light period to ensure steady state photosynthesis. Three leaves from the same plant, still attached to the mother plant, were placed in the labelling chamber ^18^ (Figure S8) and exposed to a continuous flow of a humidified, unlabelled air mixture containing: 79 % (v/v) N_2_, 21 % (v/v) O_2_, and 420 ppm ^12^CO_2_ at a flow rate of 10 L min^-1^ for 1 minute to restore steady state photosynthesis. The air mixture was generated using a stationary WMR 4008 gas mixer (Westphal Mess-und Regeltechnik GmbH, Ottobrunn, Germany) and humidified by bubbling through water in a gas wash bottle. Control samples were collected after supplying the chamber for 1 min with the unlabelled air mixture. For pulse labelling with ^13^CO_2_, the gas flow was switched to an air mixture containing: 79 % (v/v) N_2_, 21 % (v/v) O_2_, and 420 ppm ^13^CO_2_ (Figure S8A) for 10, 30 or 60 s (ME_BS&M_ x B6 lines) or for 30 and 60 s (ME_BS_ x 29 lines), before rapidly quenching the leaves with liquid N_2_ ^18,28^ (Figure S8A-B). For pulse-chase labelling, leaves were exposed to a pulse of ^13^CO_2_ for 30 s, as above, before switching back to the air mixture containing unlabelled ^12^CO_2_ (Figure S8B) for 5, 15 or 60 s (ME_BS&M_ x B6 lines) or 5, 15, 30, 60 and 120 s (ME_BS_ x 29 lines) and then rapidly quenching the leaves with liquid N_2_. The frozen leaf tissue was removed from the chamber and stored at -80°C before analysis. Plants from different lines were selected at random for labelling, and the pulse and pulse-chase treatments of different durations were performed in a randomized manner.

### [4-^13^C]malate synthesis

Enzymatic synthesis of [4-^13^C]malate was based on the protocol described in ^45^. The synthesis reaction (1 mL) contained: 10 mM HEPES-KOH pH 7.5, 13 mM phospho*enol*pyruvic acid (trisodium salt hydrate; Sigma-Aldrich, catalogue No. P7002), 10 mM MgCl_2_, 10 mM NaH^13^CO_3_ (Sigma-Aldrich, 372382), 15 mM NADH (Sigma-Aldrich, 10128023001), 4 units of PEP carboxylase (Sigma-Aldrich, C1744) and 2 units of malate dehydrogenase (Roche, 10127256001). All solutions were freshly prepared. After mixing, reaction mixture was incubated overnight at room temperature. The concentrations of the reagents were optimized to achieve quantitative conversion of NaH^13^CO_3_ into malate. The reaction was stopped by heating for 2 min at 99 °C. After cooling, the reaction mixture centrifuged at 20,000x*g* for 10 minutes to remove denatured protein and other insoluble material. Unreacted NADH was removed from solution by addition of activated charcoal, vortex mixing and incubation at room temperature for 10 minutes. The reaction mixture was then centrifuged at 20,000x*g* for 10 minutes to pellet the charcoal. The concentration of malate in the supernatant was measured enzymatically as described in ^46^ and found to be 9.8 mM. Reverse-phase high-performance liquid chromatography coupled to tandem mass spectrometry ^47^ was used to determine the isotopologue composition of the [4-^13^C]malate preparation: m_0_ (1.4%), m_1_ (98.6%), m_2_ (<0.1%), m_3_ (none detected), m_4_ (none detected). Enzymatic analysis ^48^ showed residual concentrations of 0.09 mM PEP and 0.6 mM pyruvate, the latter presumably derived from initial contamination or degradation of PEP. The synthesis reaction was optimized to ensure there was no residual NaH^13^CO_3_ in the preparation.

### Feeding ^13^C-labelled C_4_ pathway intermediates to detached rice leaves

To detect *in-vivo* fluxes at individual steps in the C_4_ pathway, detached rice leaves were supplied with ^13^C-labelled pathway intermediates inside a custom-made chamber with a controlled air flow (Figure S8C). The chamber ^49^ was constructed from a transparent Magenta™ GA-7 plant culture box (Sigma-Aldrich, catalogue No. V8505, Figure S8C) placed upside down. The box contained four holes on opposite sides fitted with transparent PVC tubing to provide two gas lines for entry and two lines for exit of a humidified air mixture, and a plastic funnel was inserted through one of the side walls of the chamber to allow liquid N_2_ to be poured into the chamber for rapid quenching of the leaves. A kneadable, solvent-free, synthetic rubber sealant (Teroson RB IX, Henkel AG & Co. KGaA, Düsseldorf, Germany) was used to make a gas-tight seal around the tubing and to close the funnel during feeding. An artificial air mixture containing: 79% (v/v) N_2_, 21% (v/v) O_2_ and either 420 ppm ^12^CO_2_ (ambient CO_2_) or 50 ppm ^12^CO_2_ (low CO_2_), was continuosly flushed through the chamber at a flow rate of 10 L min^-1^. The outlet gas line was connected to a GasHound LI-800 infrared CO_2_ analyzer (Li-Cor, Lincoln, NE, USA; https://www.licor.com/) to monitor the CO_2_ concentration, and the CO_2_ supply to the gas mixer was adjusted to maintain the desired CO_2_ concentration.

Feeding experiments were performed within the plant growth chamber between 2 to 8 hours after dawn (irradiance 560 µmol m^-2^ s^-1^, 28°C). For each sample, the apical portion of a rice leaf (approx. 10 cm from the leaf tip) was cut under water to prevent air embolism in the xylem. The cut end was placed directly in a 2 mL Eppendorf tube containing 0.2 mL of water or a solution containing: (i) 6 mM [4-^13^C]malate (see above), (ii) 6 mM [4-^13^C]aspartate (Sigma-Aldrich, catalogue No. 489999) or (iii) 2 mM [2,3-^13^C_2_]pyruvate (Sigma-Aldrich, catalogue No. 486191), and then enclosed within the feeding chamber. For feeding with mM [4-^13^C]malate or [4-^13^C]aspartate, the leaves were flushed with air mixtures containg 420 (ambient) or 50 ppm (low) CO_2_, and replicate samples were rapidly quenched with liquid N_2_ after 5, 10 or 20 min. For feeding with [2,3-^13^C_2_]pyruvate, the leaves were flushed with an air mixture containing 420 ppm CO_2_ and rapidly quenched after 5 or 10 min. After quenching, the leaf tip (∼1 cm) was excised and discarded as this segment of the leaf has a highly variable metabolite content ^50^. The remaining tissue was stored at -80°C until analysis. The 4-^13^C]malate and [4-^13^C]aspartate concentrations supplied did not increase total malate and aspartate levels in the leaves and were the highest that could be used without wilting. The [2,3-^13^C_2_]pyruvate concentration was the highest that could be used without increasing pyruvate levels in the leaf. Labelled C_4_ acids were supplied for up to 20 min but labelled pyruvate for only 5 and 10 min to minimise further metabolism and non-specific spread of label.

### Metabolite analysis

Frozen leaf material was ground to fine powder using a ball mill (Tesch, Haan, Germany) at liquid nitrogen temperature, and aliquots (15-20 mg) of frozen tissue powder were extracted with chloroform-methanol as described in ^47^. Extracts were analysed by high performance liquid chromatography coupled to tandem mass spectrometry (LC-MS/MS) to quantify isotopologues of metabolic intermediates as follows. Samples from pulse-chase labelling of the ME_BS&M_ x B6 lines lines were analyzed by reverse-phase LC-MS/MS ^47^ to measure malate, aspartate, PEP, 3PGA, dihydroxyacetone phosphate (DHAP) and 2-phosphoglycolate (2PG) and by anion-exchange LC-MS/MS ^51,52^ to measure pyruvate and citrate. Samples from pulse-chase labelling of the ME_BS_ x 29 lines lines were analyzed by reverse-phase LC-MS/MS ^47^ to measure malate, aspartate, PEP and 3PGA. Isotopologues of malate, 3PGA, F6P, FBP, PEP, pyruvate, fumarate and citrate in samples from the [4-^13^C]malate, [4-^13^C]aspartate and [2,3-^13^C_2_]pyruvate feeding experiments were analysed by anion-exchange LC-MS/MS ^51,52^. Isotopologues of aspartate, glutamate and alanine in samples from the [4-^13^C]aspartate feeding experiment and isotopologues of aspartate in samples from the [4-^13^C]malate feeding experiment were measured by hydrophilic interaction liquid chromatography coupled to tandem mass spectrometry (HILIC-MS/MS).

For the HILIC-MS/MS analysis, aliquots (10 µl) of the tissue extract were injected onto a 2.1 x 5 mm (2.7 µm particle size) Poroshell 120 HILIC-Z column (Agilent, Santa Clara, CA, USA; https://www.agilent.com) fitted with a 2.1 x 150 mm (2.7 µm) Poroshell 120 HILIC-Z guard column (Agilent), equilibrated with Eluent A (20 mM ammonium formate, pH 3, in acetonitrile). The columns were maintained at a temperature of 30°C and the flow rate was 0.45 mL min^-1^). Amino acids were eluted by mixing Eluent A and Eluent B (20 mM ammonium formate, pH 3, in water) to generate the following gradient: 0-11.5 min, 100-70% A; 11.5-13.5 min, 70% A (isocratic); 13.5-14 min, 70-100% A; 14-20 min, 100%& A (isocratic). The eluate passed directly into a Q-Trap 5500 (SCIEX, Toronto, Canada; https://sciex.com) operated in multiple reaction monitoring mode using electrospray ionization source in either negative mode (for aspartate and glutamate) or positive mode (all other amino acids). The ion transfer tube temperature was kept at 330 °C, with a capillary voltage of 1500 V. For each amino acid isotopologue, the retention time (RT) and the parent and product ion *m*/*z* settings for the first (Q1) and third (Q3) quadrupoles are shown in Supplementary Dataset 1J, along with the declustering potential (DP), entrance potential (EP), collision energy (CE) and cell exit potential (CXP) settings.

For all of the LC-MS/MS analyses, calibration curves were generated using authentic standards. Total amounts were calculated by summing isotopologue abundances. Data were corrected for natural abundance of ^2^H, ^13^C and ^15^N using the CORRECTOR software tool (https://www.mpimp-golm.mpg.de/719693/Bioinformatik-Tools).

### Statistical Analysis

Statistical analyses were performed with GraphPad Prism 10 using one-way ANOVA with Tukey’s post-hoc testing.

## ACKNOWLEDGEMENTS

We are grateful to Julian Hibberd and Asaph Cousins for careful and critical reading of the manuscript.

## AUTHOR CONTRIBUTIONS

JEL, MS, SA, CB, RTF & JAL designed research; ME generated and characterized the new transgenic rice lines; FD & KY performed crosses and characterized the progeny; SvC & RTF supervised ME, FD and KY (Figure 2; Figure S1; Table S1); SA, CB, RF, HI & JEL performed pulse chase and feeding experiments (Figures 3-5, Figures S2-S8, Table S2); SA, CB, JEL & MS analysed data (Supplementary Data, Supplementary Calculations); SA, CB, MS, JEL, RTF & JAL wrote the paper. All authors read and edited the manuscript prior to submission.

## SUPPLEMENTARY INFORMATION

### Extended data file

**Supplementary Figure 1.** Generation of transgenic lines with high levels of maize NADP-ME.

**Supplementary Figure 2.** Isotopologue distribution in the pulse.

**Supplementary Figure 3.** Isotopologue distribution in the chase.

**Supplementary Figure 4.** Correction of m*_1_* malate and m*_1_* aspartate for continued synthesis in the chase from m*_1_* PEP.

**Supplementary Figure 5.** Statistical analyses, isotopologue distribution and enrichment in selected metabolites after [4-^13^C] malate-feeding at 420 and 50 ppm CO_2_

**Supplementary Figure 6.** Statistical analysis, isotopologue distribution and enrichment in selected metabolites after [4-^13^C] aspartate-feeding at 420 and 50 ppm CO_2_.

**Supplementary Figure 7.** Statistical analysis and isotopologue distribution after [2,3-^13^C_2_] pyruvate-feeding.

**Supplementary Figure 8.** Set up for ^13^CO_2_ pulse - ^12^CO_2_ chase labelling and feeding experiments.

**Supplementary Table 1**. *In vitro* enzyme activities in transgenic rice lines: absolute rates, and as percentage of in vitro activities in maize.

**Supplementary Table 2.** Data analysis to decrease experimental noise by utilizing co-variation between labelling of different metabolites

### Datasets

**Supplementary datasets.xlxs**

Sheets A-C: Pulse chase;

Sheets D-E: [4-^13^C]malate-feeding;

Sheets F-G: [4-^13^C]aspartate-feeding;

Sheets H-I: [2,3-^13^C_2_]pyruvate-feeding experiments;

Sheet J: HILIC-MS/MS MRM settings.

### Calculations

**Supplementary calculations.xlxs**

