## Supplementary Figures S1-S8, Supplementary Tables S1-S2 for "C_4_ photosynthetic pathway fluxes in transgenic rice plants"

**Baccolini et al.**

### In the Extended data file


| *Data displays* | | *Page* |
|---|---|---|
| **Supplementary Figure S1** | Generation of transgenic lines with high levels of maize NADP-ME | 2 |
| **Supplementary Figure S2** | Isotopologue distribution in the pulse. | 3 |
| **Supplementary Figure S3** | Isotopologue distribution in the chase. | 4 |
| **Supplementary Figure S4** | Correction of $m_1$ malate and $m_1$ aspartate for continued synthesis in the chase from $m_1$ PEP. | 5-6 |
| **Supplementary Figure S5.** | Statistical analyses, isotopologue distribution and enrichment in selected metabolites after [4-$^{13}$C] malate-feeding at 420 and 50 ppm $CO_2$ | 7-8 |
| **Supplementary Figure S6** | Statistical analysis, isotopologue distribution and enrichment in selected metabolites after [4-$^{13}$C] aspartate-feeding at 420 and 50 ppm $CO_2$. | 9-10 |
| **Supplementary Figure S7**. | Statistical analysis and isotopologue distribution after [2,3-$^{13}C_2$] pyruvate-feeding. | 11-12 |
| **Supplementary Figure S8**. | Set up for $^{13}CO_2$ pulse -$^{12}CO_2$ chase labelling and feeding experiments | 13-14 |
| **Supplementary Table S1** | *In vitro* enzyme activities in transgenic rice lines: absolute rates, and as percentage of in vitro activities in maize | 15 |
| **Supplementary Table S2** | Data analysis to decrease experimental noise by utilizing co-variation between labelling of metabolites to estimate relative fluxes. | 16 |


### As separate files

**Supplementary dataset file.xlxs**

Sheets A-C: Pulse chase;

Sheets D-E: [4-$^{13}$C]malate-feeding;

Sheets F-G: [4-$^{13}$C]aspartate-feeding;

Sheets H-I: [2,3-$^{13}C_2$]pyruvate-feeding experiments;

Sheets J: HILIC-MS/MS MRM settings.

**Supplementary calculations file.xlxs**

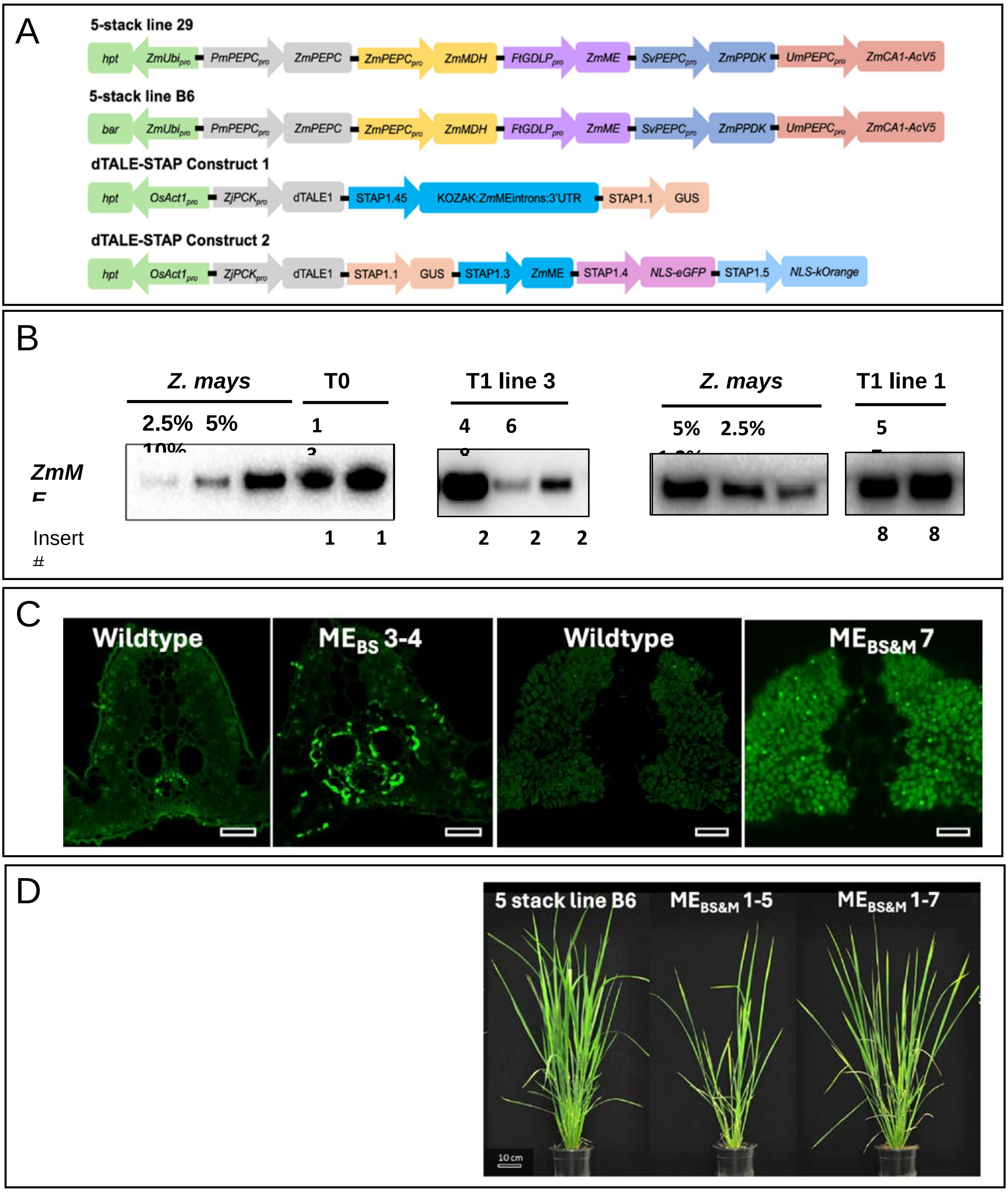


**Supplementary Figure 1. Generation of transgenic rice lines with high levels of maize NADP-ME**. **A)** Schematic of constructs in parental lines 29 and B6 [18] and in the new dTALE-STAP ME lines. **B)** Western blots showing ME levels in T0 and T1 dTALE-STAP lines. T1 line 3 contained construct 1 and T1 line 1 contained construct 2. **C)** Immunolocalization showing bundle sheath cell localization of ME in a T1 line containing construct 1 ($ME_{BS}$3-4) and in both cell-types in a T0 plant containing construct 2 ($ME_{BS\&M}$-7) (referred to collectively as 'high-ME lines'). **D)** Phenotype of T1 high-ME plants selected as parents for crosses. $ME_{BS}$3-4 was the parent for both the $ME_{BS}$ x 29-A-6-3 and $ME_{BS}$ x 29-C-1-5 lines. $ME_{BS\&M}$1-5 was the parent for the $ME_{BS\&M}$ x B6-A-5 line and $ME_{BS\&M}$1-7 was the parent for the $ME_{BS\&M}$ x B6-B line.

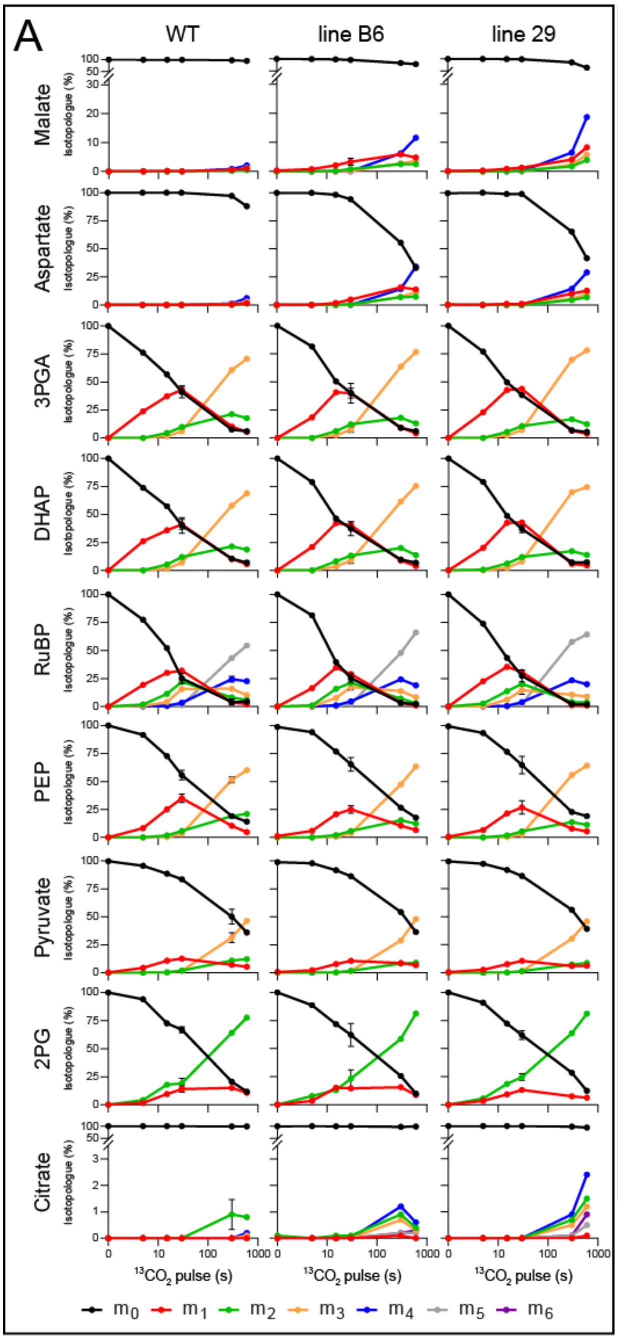


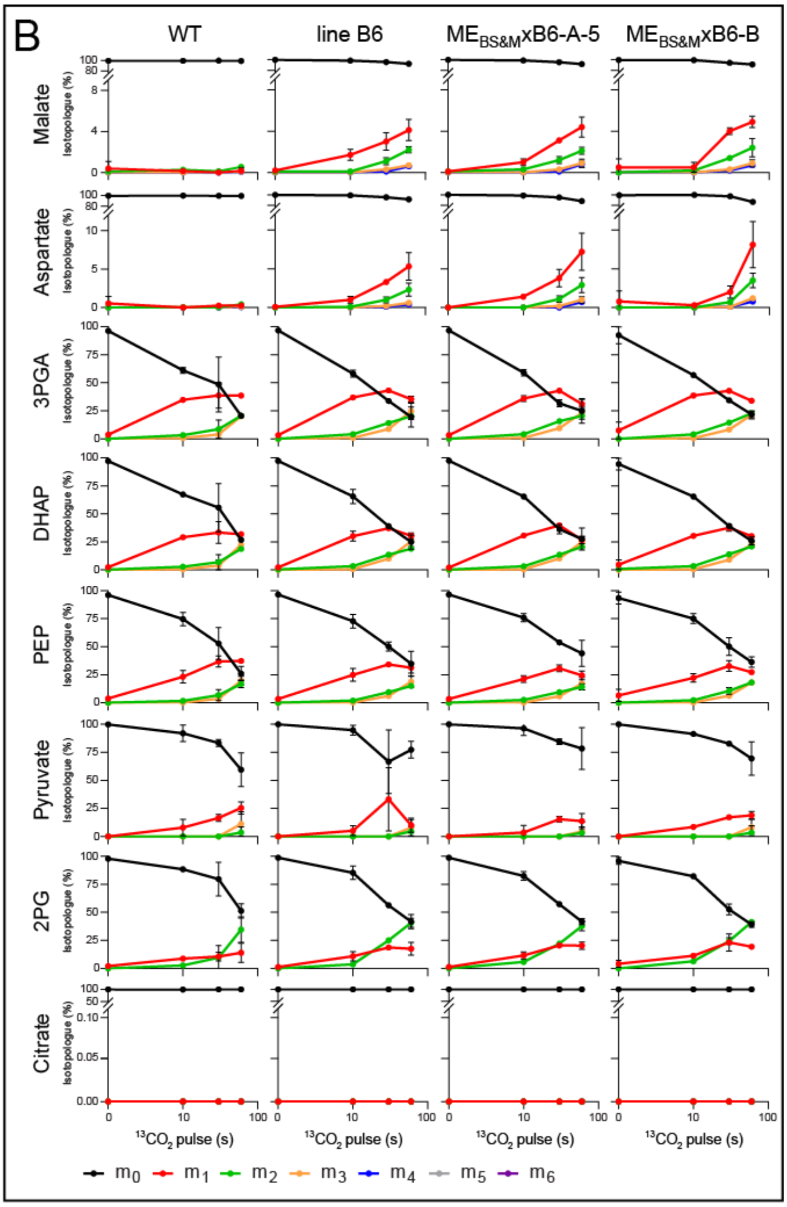


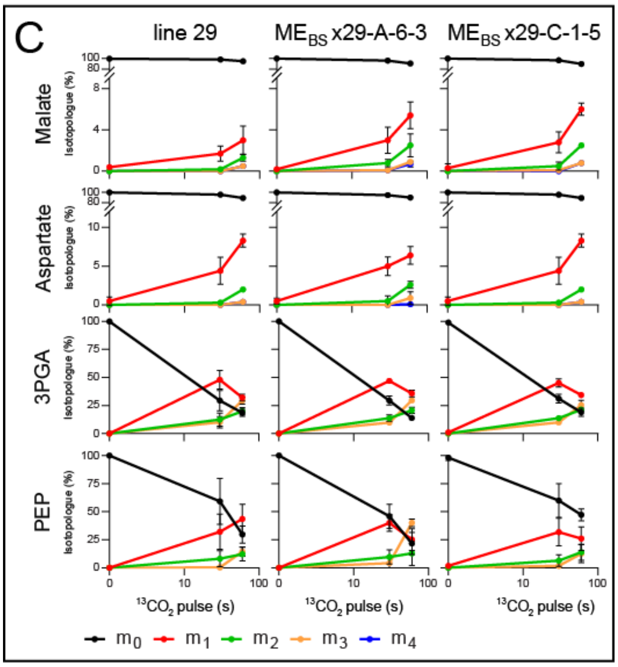


**Supplementary Figure 2.** Isotopologue distribution in the pulse. The relative abundance of each isotopologue ($m_n$) for a given metabolite is represented, where $_n$ is the number of $^{13}$C atoms present in the molecule. The x-axes correspond to the labelling time on a $log_{10}$ scale. A y-axis break is used to improve visualization of labelled isotopologues for metabolites with low labelling. The pulse experiments were performed in parallel with the chase experiments in Figure S3. Note that the maximum pulse time was longer in the experiment from Ermakova *et al* (2021) [18] with the parental lines (panel A) than with the $ME_{BS\&M}$x B6 (panel B) and $ME_{BS}$ x 29 (panel C) experiments. Accumulation of labelled isotopologues was estimated from linear regression of time from 0-600 s for wild-type and 0-30 s for lie B6 and line 29 in panel A, and from 0-60 s for all genotypes in panels B and C. Mean ± SD, n = 1 to 4, 2 to 3 and 3 to 7 for each labelling experiment. The raw data are provided in Supplementary Datasets S1A-C.

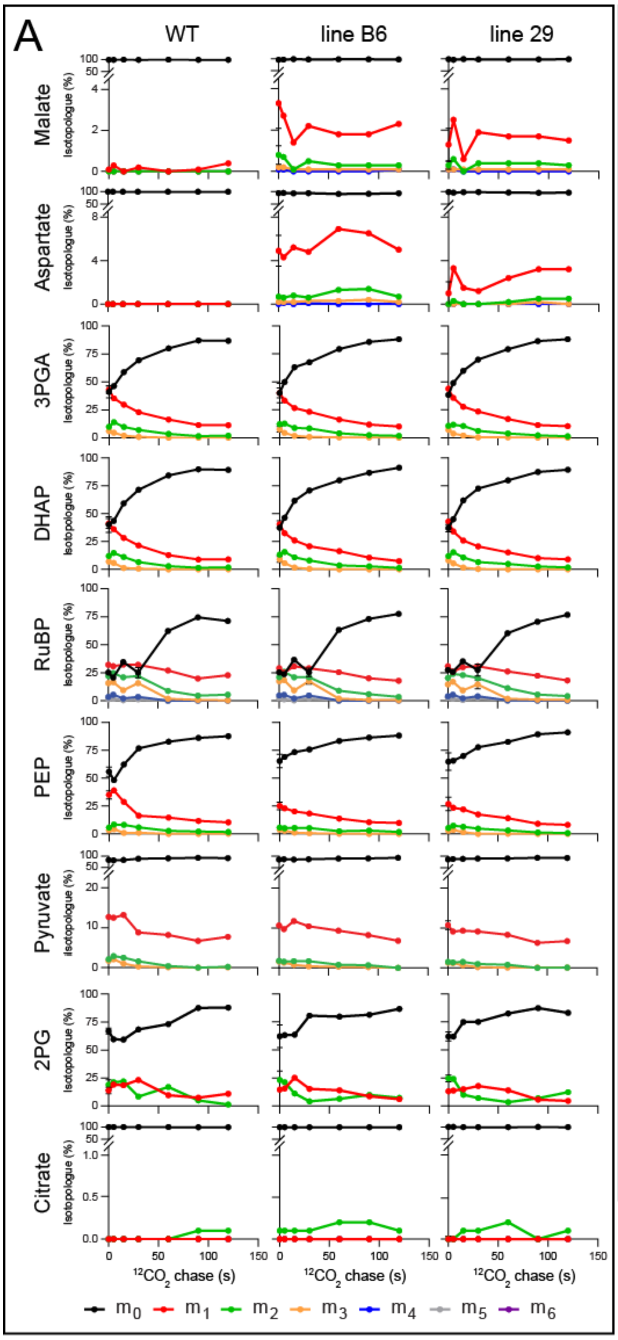


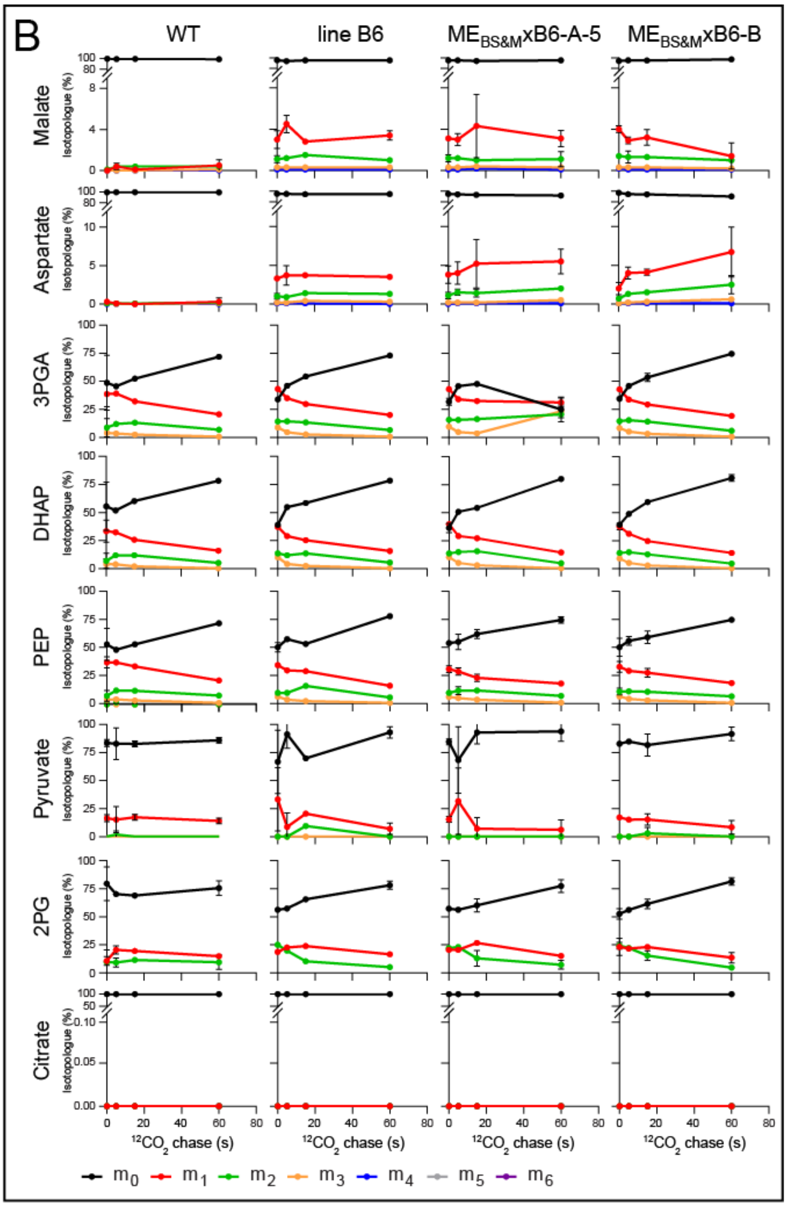


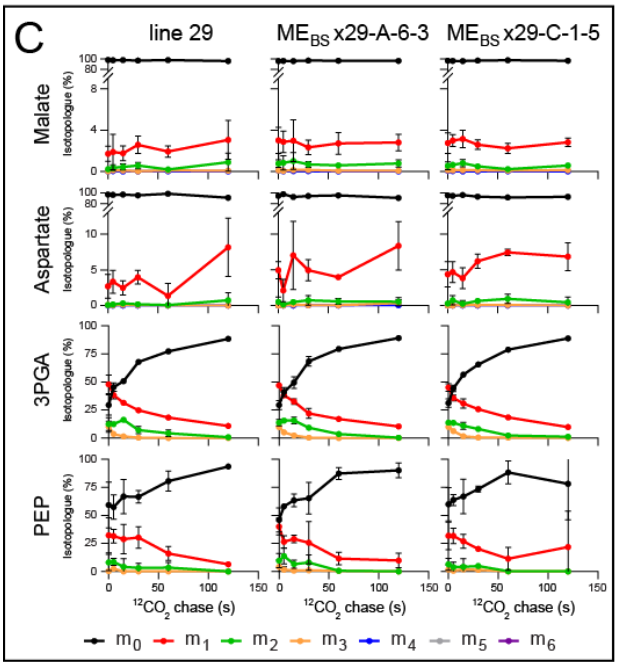


**Supplementary Figure 3.** Isotopologue distribution in the chase. The relative abundance of each isotopologue ($m_n$) for a given metabolite is represented, where $_n$ is the number of $^{13}C$ atoms present in the metabolite. The x-axes correspond to the labelling time (linear scale). A y-axis break is used to improve visualization of labelled isotopologues for metabolites with low labelling. The chase experiments were performed in parallel with the pulse experiments (Figure S2). In all cases, the preceding pulse was 30 sec. The duration of the chase was 120 sec in Ermakova *et al*. (2021) [18] (panel A) and the $ME_{BS}$ x 29 experiment (panel C), and 60 sec in the $ME_{BS\&M}$x B6 experiment (panel B). Mean ± SD, n = 1 to 4, 1 to 3 and 3 to 6 for each labelling experiment. The raw data are provided in Supplementary Datasets 1A-C.

**Supplementary Figure 4. Correction of $m_1$ malate and $m_1$ aspartate for continued synthesis in the chase from $m_1$ PEP. A)** Plots showing measured abundance of $m_1$ malate, corrected abundance of $m_1$ malate, measured abundance of $m_1$ aspartate and corrected abundance of $m_1$ aspartate for the chase in the three pulse-chase experiments with parental lines (from Ermakova *et al.* [18], $ME_{BS\&M}$x B6 and $ME_{BS}$ x 29 lines. For the correction of the malate data, the average fractional abundance of the $m_1$ isotopologue of PEP up to a given time, $t_x$, in the chase was multiplied by the rate of conversion of PEP to malate (estimated as the rate of accumulation of summed labelled isotopologues of malate in the pulse - see Supplementary Calculations) to estimate the amount of $m_1$ malate synthesized from $m_1$ PEP between the start of the chase and time $t_x$ in the chase. This value was subtracted from the measured abundance of $m_1$ malate at time $t_x$ to provide a corrected value, which estimates how much of the $m_1$ malate that was synthesized in the pulse remained at time $t_x$ in the chase. This procedure was performed for each time point in the chase. An analogous procedure was followed for aspartate. The size of the correction can be seen by comparing the measured $m_1$ abundance and corrected $m_1$ abundance plots. In some cases, the corrected value is negative because more $m_1$ isotopologue was synthesized from $m_1$ PEP in the chase than was synthesized from $^{13}CO_2$ in the short pulse. The plots show each individual sample. Linear regression and fitted exponential decay lines are presented as blue and red lines, respectively. Note that the y-axis scale is selected to show the full range of the data points and varies between genotypes (in particular, the scale is greatly expanded for wild-type, WT) and whether $m_1$ malate or $m_1$ aspartate is shown. **B)** The summed absolute amount of $m_1$ malate and $m_1$ aspartate (nmol $m_1$ $g^{-1}$FW), after correction for the amount of $m_1$ isotopologue synthesized from $m_1$ PEP in the chase. The slope of the regression indicates the rate of $^{13}C$ release by decarboxylation (nmol $^{13}C$ $g^{-1}$ FW $s^{-1}$). Note that the scale of the y-axis differs depending on the genotype, and that the chase was for up to 120 sec in Ermakova *et al.* [1] and for the $ME_{BS}$ x 29 lines, and 60 sec for the $ME_{BS\&M}$x B6 lines. The slopes, $R^2$ and p-values are summarized in Figure 3E. **C)** Initial decay rates estimated by fitting an exponential decay curve. The chase data for corrected $m_1$ malate abundance, corrected $m_1$ aspartate abundance, and the combined abundance of corrected $m_1$ malate and aspartate were fitted with an exponential decay function $y=y_0 \cdot e^{-k \cdot x}$, where k represents the rate constant. Errors on the slopes were determined by propagating the uncertainties on k and $y_0$. The goodness of fit is expressed as $R^2$. n.d., not determined, indicates cases where the selected model did not adequately describe the data. The slopes estimated from a linear regression on combined corrected $m_1$ malate and corrected $m_1$ aspartate are shown for comparison (taken from Figure 3E). Significant slopes ($p < 0.05$) for linear regression are highlighted in bold. Mal = malate, Asp = aspartate. See Supplementary Calculations for raw data.

Figure overpage

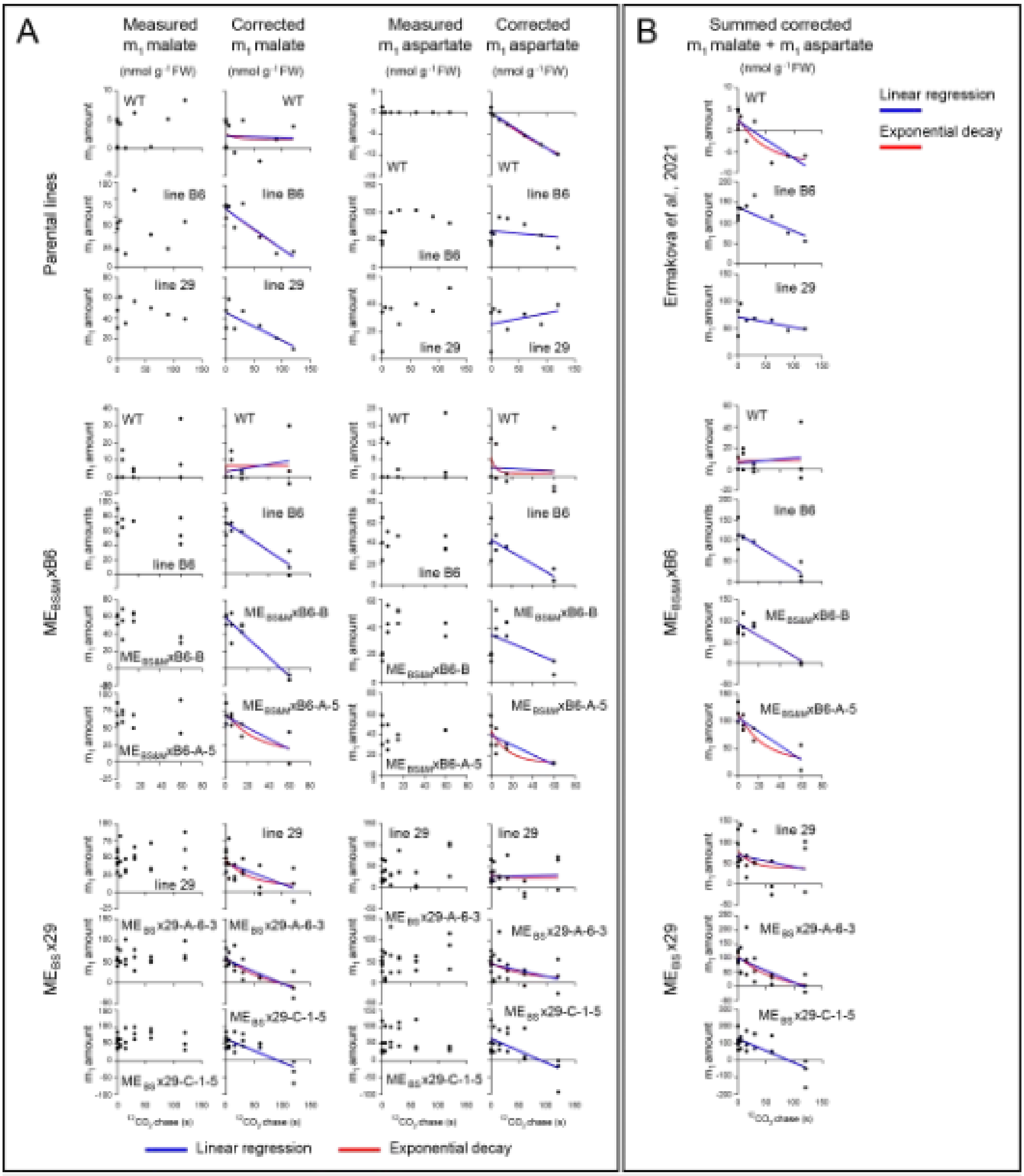


C

| | Line | Estimated by fitting an exponentional decay | | | | | | | | | Linear regression | | |
|---|---|---|---|---|---|---|---|---|---|---|---|---|---|
| | | Corrected $m_1$ malate (nmol $g^{-1}$ FW $s^{-1}$) | | | Corrected $m_1$ Asp (nmol $g^{-1}$ FW $s^{-1}$) | | | Corrected $m_1$ Mal+Asp (nmol $g^{-1}$ FW $s^{-1}$) | | | Corrected $m_1$ Mal+Asp (nmol $g^{-1}$ FW $s^{-1}$) | | |
| | | slope | error | $R^2$ | slope | error | $R^2$ | slope | error | $R^2$ | slope | $p$ | $R^2$ |
| Parental lines | WT | -0.01 | ± 0.03 | 0.005 | n.d | n.d. | -0.568 | -0.40 | ± 1.77 | 0.166 | -0.09 | **0.004** | 0.67 |
| | line B6 | -0.78 | ± 0.29 | 0.766 | -0.09 | ± 0.20 | 0.038 | -0.67 | ± 0.35 | 0.512 | -0.58 | **0.019** | 0.57 |
| | line 29 | -0.37 | ± 0.22 | 0.560 | 0.00 | ± 0.11 | -0.094 | -0.22 | ± 0.23 | 0.193 | -0.19 | 0.269 | 0.20 |
| $ME_{BS4M}$xB6 | WT | 0.00 | ± 0.14 | -0.061 | -1.01 | ± 2.55 | 0.066 | 0.00 | ± 0.20 | -0.019 | 0.00 | 0.653 | 0.02 |
| | line B6 | -1.95 | ± 0.82 | 0.807 | -1.13 | ± 0.68 | 0.664 | -3.07 | ± 1.41 | 0.776 | -1.59 | **0.001** | 0.79 |
| | $ME_{BS4M}$xB6-B | -2.13 | ± 1.14 | 0.738 | -0.34 | ± 0.38 | 0.186 | -2.11 | ± 1.18 | 0.635 | -1.46 | **0.000** | 0.77 |
| | $ME_{BS4M}$xB6-A-5 | -1.54 | ± 0.79 | 0.637 | -1.01 | ± 0.57 | 0.589 | -2.54 | ± 0.94 | 0.771 | -1.28 | **0.002** | 0.76 |
| $ME_{BS}$x29 | line 29 | -0.73 | ± 0.40 | 0.395 | 0.00 | ± 0.15 | -0.002 | -0.54 | ± 0.53 | 0.087 | -0.28 | 0.256 | 0.07 |
| | $ME_{BS}$x29-A-6-3 | -1.43 | ± 0.70 | 0.598 | -0.60 | ± 0.57 | 0.160 | -2.03 | ± 1.15 | 0.454 | -0.83 | **0.001** | 0.43 |
| | $ME_{BS}$x29-C-1-5 | -0.91 | ± 0.63 | 0.417 | -0.95 | ± 0.95 | 0.266 | -1.86 | ± 1.49 | 0.356 | -1.41 | **0.000** | 0.52 |

6

Supplementary Figure S4

**Supplementary Figure 5. Statistical analyses, isotopologue distribution and enrichment in selected metabolites after [4-$^{13}$C]malate-feeding at 420 and 50 ppm $CO_2$.** A) Relative abundance of isotopologues for malate, aspartate and 3PGA. Values are mean ± SD (n = 3 to 6). **B)** Statistical analyses for individual metabolites, for each feeding duration and $CO_2$ amount. Analyses are provided for the key parameters: (i) fractional abundance of $m_1$ malate and $m_1$ aspartate (malate $m_1$ %, aspartate $m_1$ %) that provide information about label intensity in the C4 position of the substrate for decarboxylation, and (ii) 3PGA % enrichment that provides information about release of $^{13}CO_2$ by decarboxylation and reassimilation by Rubisco. Enrichment is also shown for: (iii) metabolites downstream of 3PGA (FBP, F6P PEP) that provide further support for flow of $^{13}$C into the CBC, and (iv) organic acids (pyruvate, fumarate, citrate) that provide information about further metabolism of the introduced [4-$^{13}$C]malate. The plots shows the individual data points and the mean ± SD (n = 6 for wild-type and 3 for the transgenic lines). Statistical analyses were performed using one-way ANOVA with Tukey's post-test. Only statistically significant differences are displayed and are indicated (* for $p < 0.05$, ** for $p < 0.01$, *** for $p < 0.005$, **** for $p < 0.001$). Note that for a given metabolite, to display the full range of the data, different y-axes were used for each feeding duration and for $CO_2$ amounts. **C)** $^{13}$C enrichment in FBP, F6P, PEP, pyruvate, fumarate and citrate. Values are mean ± SD (n = 3 to 6). Original data is shown in Supplementary Datasets S1D-E.

Figure overpage

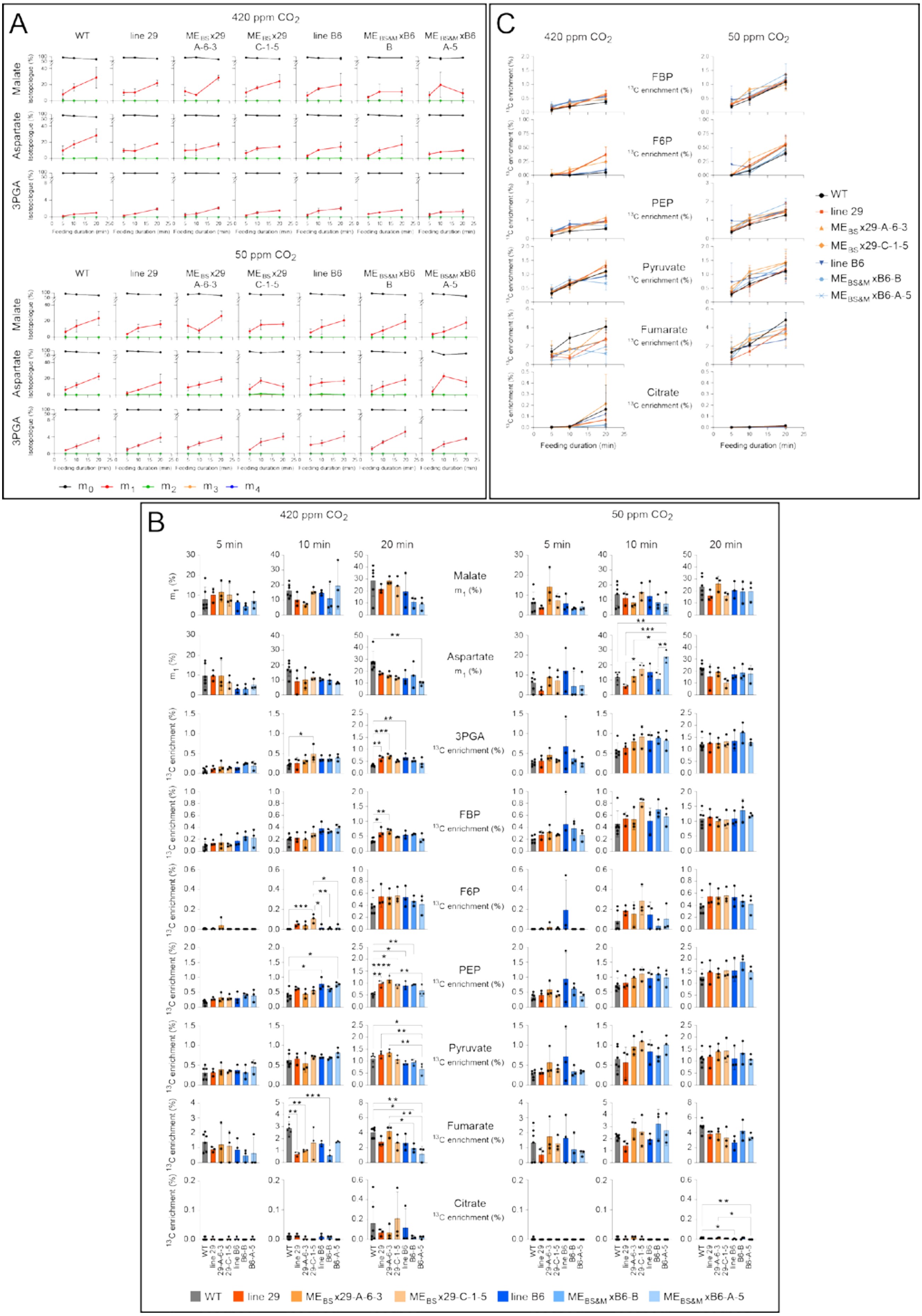


Supplementary Figure S5

**Supplementary Figure 6. Statistical analyses, isotopologue distribution and enrichment in selected metabolites after [4-$^{13}$C]aspartate-feeding at 420 and 50 ppm $CO_2$. A)** Relative abundance of isotopologues for aspartate, malate and 3PGA. Values are mean ± SD (n = 3 to 6). **B)** Statistical analyses for individual metabolites, for each feeding duration and $CO_2$ amount. Analyses are provided for the key parameters: (i) fractional abundance of $m_1$ aspartate and $m_1$ malate (aspartate $m_1$ %, malate $m_1$ %) that provide information about label intensity in the C4 position of the substrate for decarboxylation, and (ii) 3PGA % enrichment that provides information about release of $^{13}CO_2$ by decarboxylation and reassimilation by Rubisco. Enrichment is also shown for: (iii) metabolites downstream of 3PGA (FBP, F6P PEP) that provide further support for flow of $^{13}$C into the CBC, and (iv) organic acids (pyruvate, fumarate, citrate) that provide information about further metabolism of the introduced [4-$^{13}$C]malate. The display shows the individual data points and the mean ± SD (n = 6 for wild-type and 3 for the transgenic lines). Statistical analyses were performed using one-way ANOVA with Tukey's post-test. Only statistically significant differences are displayed and are indicated (* for $p < 0.05$, ** for $p < 0.01$, *** for $p < 0.005$, **** for $p < 0.001$). Note that for a given metabolite, to display the full range of the data, different y-axes were used for each feeding duration and for $CO_2$ amounts. **C)** $^{13}$C enrichment in FBP, F6P, PEP, pyruvate, fumarate and citrate. Values are mean ± SD (n = 3 to 6). Original data is in Supplementary Datasets S1F-G.

Figure overpage

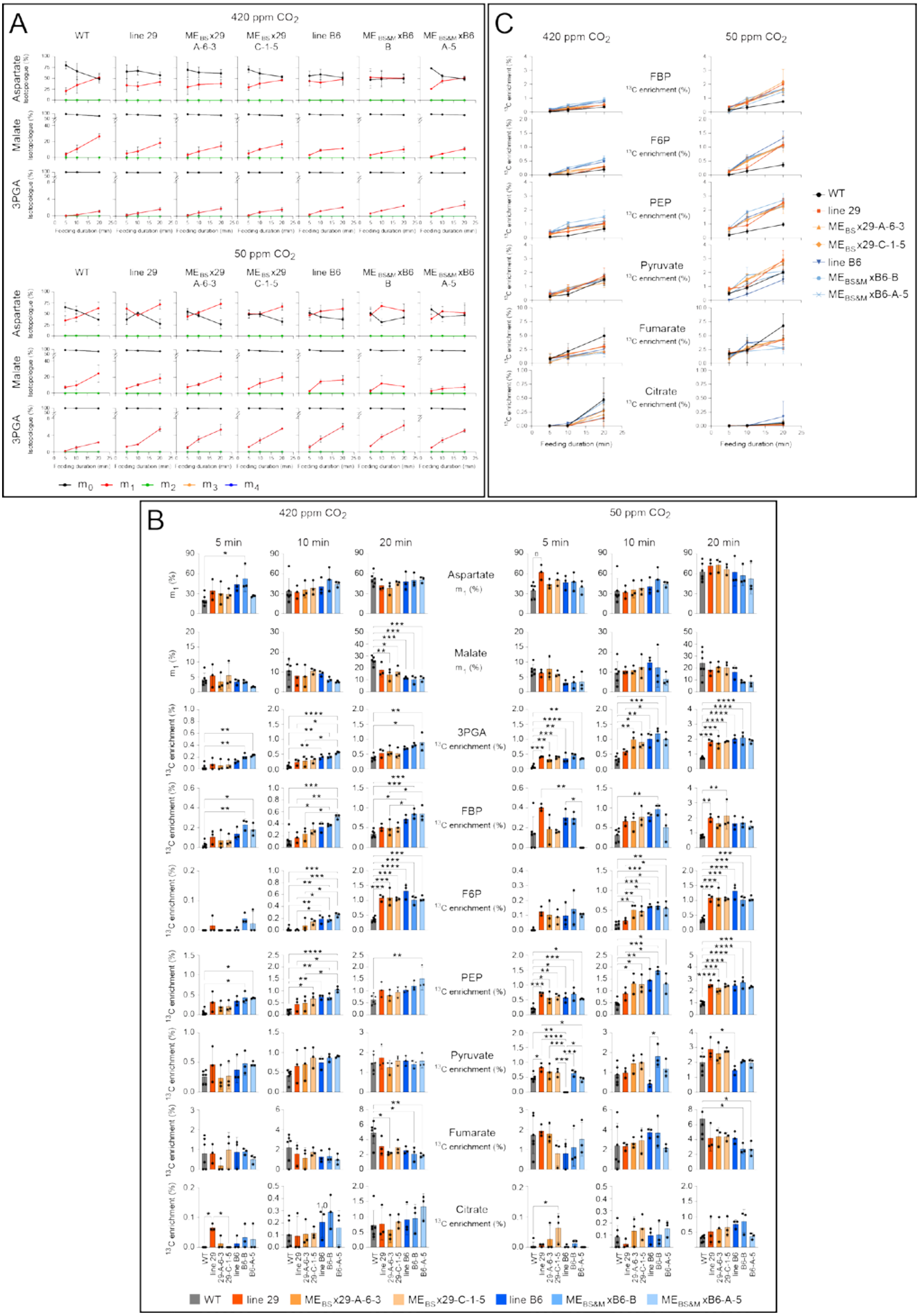


Supplementary Figure S6

**Supplementary Figure 7. Statistical analysis and isotopologue distribution after [2,3-$^{13}C_2$]pyruvate-feeding. A)** Relative abundance of isotopologues over feeding duration. Plots show mean ± SD (n = 4 for wild-type and 3-4 for transgenic lines). **B)** Statistical analyses for individual metabolites, for each feeding duration and experiment. Analyses are shown for key parameters: (i) fractional abundance of $m_2$ pyruvate (% $m_2$ pyruvate) that provides information about label intensity in the C2 and C3 position of the substrate for PPDK and (ii) fractional abundance of $m_2$ PEP (% $m_2$ pyruvate) that provides information about flow of label via PPDK. Analyses are also shown for fractional abundance of $m_2$ (% $m_2$) in: (iii) metabolites downstream of PEP (3PGA, FBP, F6P) and (iv) organic acids (malate, citrate, fumarate) that provide information about the extent to which [2,3-$^{13}C_2$]pyruvate was metabolized by other routes. Note that for given metabolite, different y-axes were used for each feeding duration and experiments to show the full range of the data. The display shows the individual data points and the mean ± SD (n = 3 - 6). Statistical analyses were performed using one-way ANOVA with Tukey's post-test. Only statistically significant differences are displayed and are indicated (* for $p < 0.05$, ** for $p < 0.01$, *** for $p < 0.005$, **** for $p < 0.001$). Note that for a given metabolite, different y-axes were used for each feeding duration and experiment to show the full range of the data. Original data is in Supplementary Datasets S1H-I.

Figure overpage

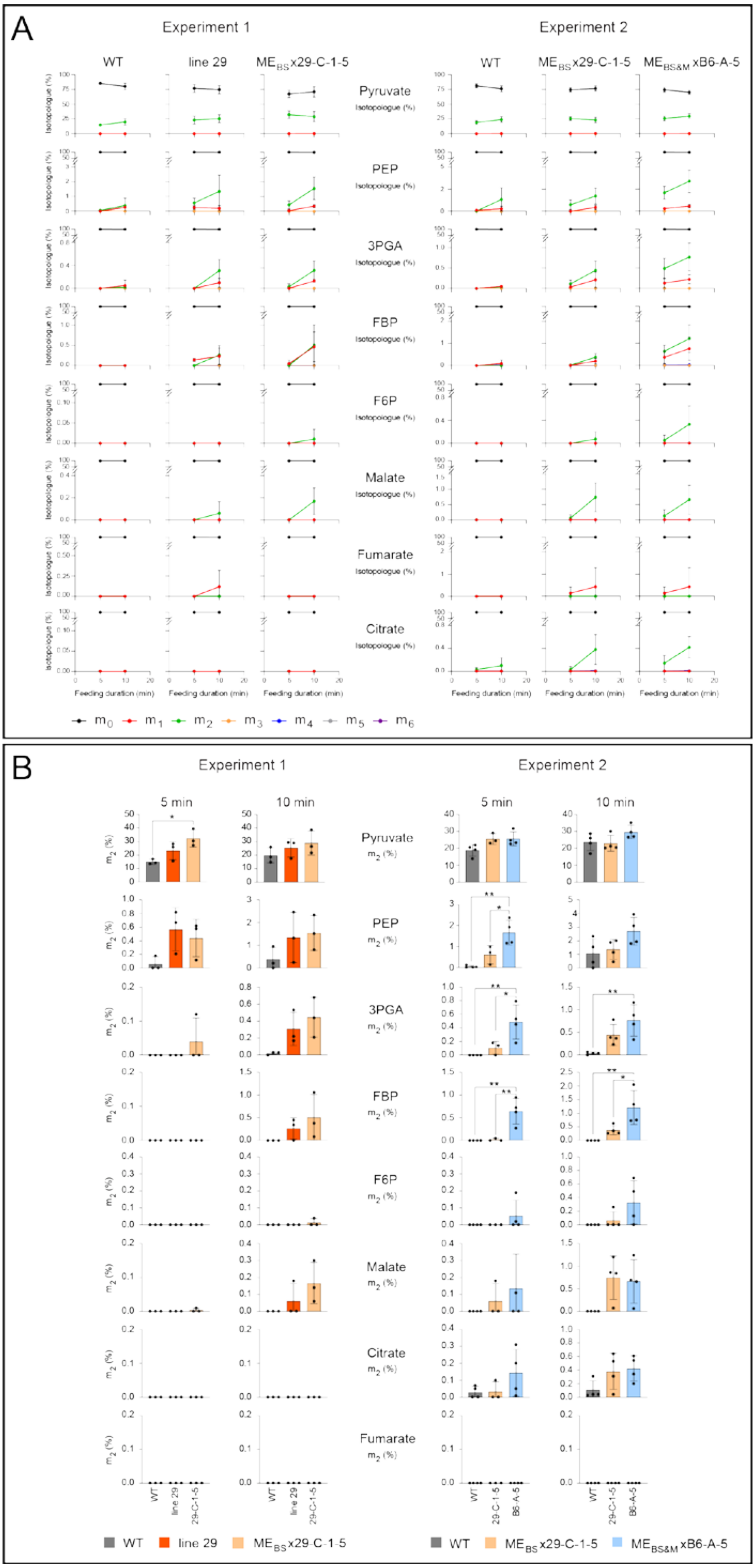


Supplementary Figure S7

S**upplementary Figure 8. Set up for $^{13}CO_2$ pulse – $^{12}CO_2$ chase labelling and feeding experiments.** Rice plants were grown in a growth chamber with a 14-h photoperiod, an irradiance of 560 µmol $m^{-2}$ $s^{-1}$ (provided by a mixture of white and far-red LEDs tuned to mimic the spectrum of sunlight), day/night temperatures of 28°C/20°C and a relative humidity of 80%. Plants were labelled *in situ* within the growth chamber to maintain steady state rates of photosynthesis. **A, B)** $^{13}CO_2$ pulse-chase labelling. Two gas mixers (light grey rectangle) generate two synthetic air mixtures: one comprising 79% $N_2$, 21% $O_2$ and 420 ppm $^{12}CO_2$ (blue line) and another one containing 79% $N_2$ and 21% $O_2$ (green line). These gas mixtures are directed into the plant growth chamber, where they are humidified by bubbling through water (28°C). A separate commercially purchased gas cylinder supplies $^{13}CO_2$ and is connected (not shown) to a mass flow controller that regulates flow. The $^{13}CO_2$ (red line) is mixed with the 79% $N_2$ and 21% $O_2$ mixture using a T-shaped gas mixer to provide a $^{13}CO_2$ concentration of 420 ppm in the $^{13}CO_2/N_2/O_2$ mixture. A 4–way ball valve controls which gas mixture enters the labelling chamber. During a $^{13}CO_2$ pulse (A) it directs the air mixture containing $^{13}CO_2$ towards the labelling chamber and the $^{12}CO_2$-containing air mixture towards a soda lime $CO_2$ trap. The gas outlet port of the labelling chamber is connected to a $CO_2$ trap located outside the phytotron. The $CO_2$ traps serve to prevent $^{13}CO_2$ release into the growth chamber, as this might result in $^{13}CO_2$ uptake by other plants and compromise later experiments by increasing the background abundance. In the chase (B), the opposite configuration is used. Pre-mixing of these two gas mixtures and use of a 4-way ball valve is essential to allow instantaneous switching from the $/N_2/O_2/^{12}CO_2$ mixture to the $N_2/O_2/^{13}CO_2$ mixture at the start of the pulse, and from the $N_2/O_2/^{13}CO_2$ mixture to the $N_2/O_2/^{12}CO_2$ mixture at the start of the chase. To minimise the delay in changing the gas composition in the labelling chamber, the tube between the T-shaped mixer and the labelling chamber and the chamber volume are minimised. Chamber volume is 83.5 ml and flow rate 10 L $min^{-1}$, giving a halftime of 0.35 s for renewal of gas in the chamber. Rapid quenching of leaf metabolism is achieved by pouring liquid nitrogen (equivalent to several times the internal volume of the labelling chamber) through a funnel connected to the entry port of the labeling chamber. An exit port, situated diagonally opposite the entry port, allows $N_2$ to flood and then flow out of the chamber. During labelling, the entry and exit ports are sealed with Teflon plugs (not shown). For each labelling time point, three attached leaf blades (mid sections) were clamped in the labelling chamber and flushed with the air mixture containing $^{12}CO_2$ for 1 min, to restore steady state photosynthesis before labelling commenced. **C**) Feeding set up. A gas mixer generates a synthetic air mixture containing 79% $N_2$, 21% $O_2$, and 420 or 50 ppm $^{12}CO_2$ (blue line), which is humidified by bubbling through water at 28°C. The gas line is then split into four lines using three Y-shaped hose connectors (dark blue). This configuration allows two feeding boxes to operate in parallel, with each box receiving gas through two dedicated lines. The feeding boxes are equipped with a funnel through which liquid nitrogen is poured to quench metabolism and a gas outlet line. During the feeding, the funnel is closed by a plug made with Blu Tack® putty (not shown). The outlet gas line is connected to an infrared gas analyzer and to a soda-lime $CO_2$ trap. For each feeding time point, the distal part of a rice leaf blade was cut under water, placed in a 2-mL Eppendorf tube containing either 0.2 mL of feeding solution or water, and positioned inside the labelling box. Detached leaves were supplied labelled malate, aspartate or pyruvate *in situ* within the growth chamber. Figure adapted from [56].

Figure overpage

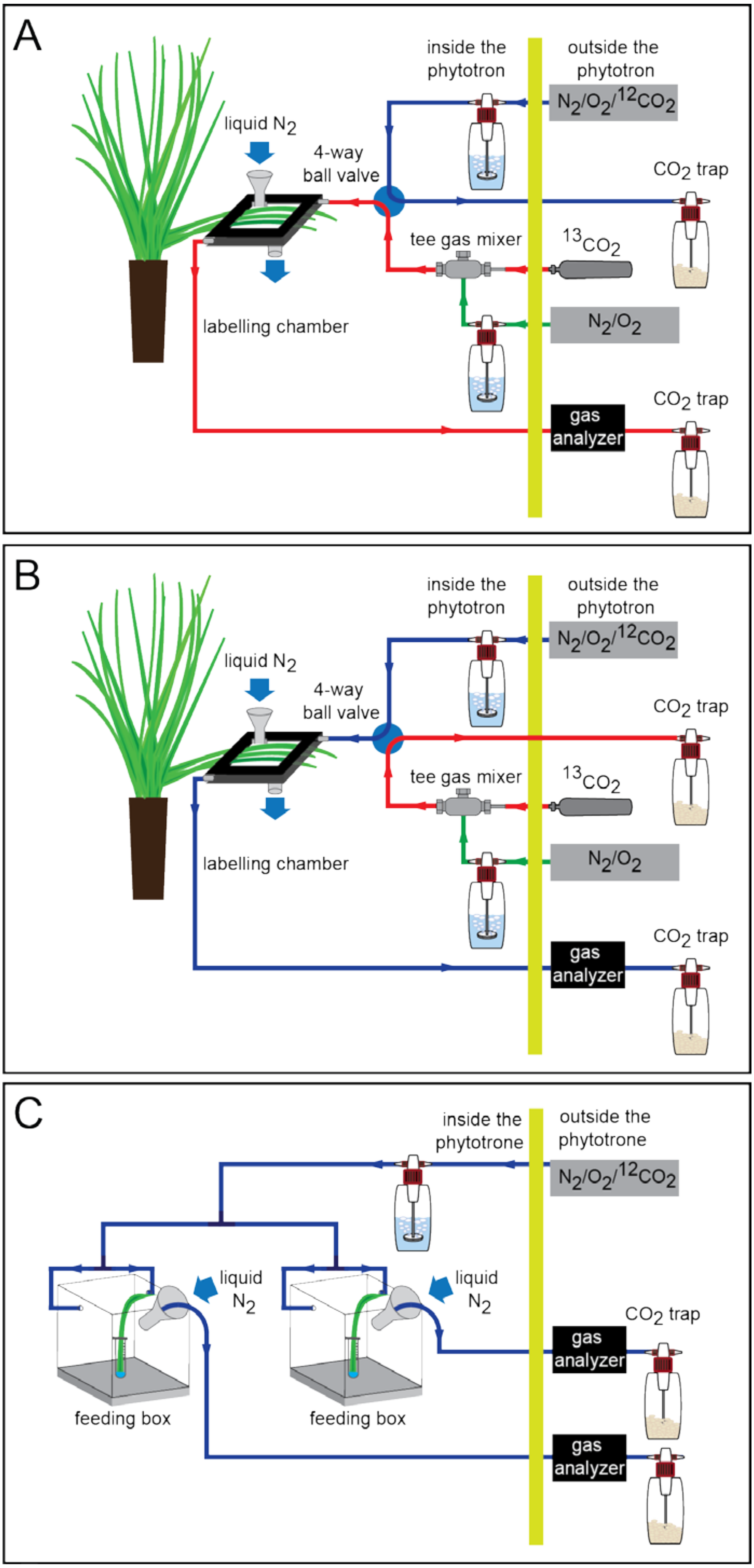


Supplementary Figure S8

| Genotype | Extractable Activity (nmol $g^{-1}$ FW $s^{-1}$) | | |
|---|---|---|---|
| | **PEPC** | **NADP-ME** | **PPDK** |
| Wild-type | 18.2±1.7 (1.1%) | 1.1±0.2 (0.4%) | 2.3±0.8 (1.2%) |
| Line B6 | 54.1±4.9 (3.4%) | 7.3±0.4 (3%) | 26.7±2.0 (14%) |
| $ME_{BS\&M}$ x B6-A-5 | | 38.3±9.1 (16%) | |
| $ME_{BS\&M}$ x B6-B | | 28.1±6.0 (12%) | |
| Line 29 | 53.0±7.7 (3.4%) | 4.5±0.2 (2%) | 27.7±1.1 (15%) |
| $ME_{BS}$ x 29-A-6-3 | | 72.3±3.5 (30%) | |
| $ME_{BS}$ x 29-C-1-5 | | 92.3±6.0 (38%) | |

**Supplementary Table 1. *In vitro* enzyme activities in transgenic rice lines:** absolute rates, and as percentage of *in vitro* activities in maize

Maximum extractable activities of NADP-ME from wild type rice, transgenic parents Line 29 and Line B6 (described in [1]), and crosses used for labelling expressed on a fresh weight basis (n=4). Activities of PEPC and PPDK in the parental lines are also shown. Values in brackets are % of maize activities reported in Ermakova *et al* (2021) [18].

| LINE | Malate feeding, 420 ppm $CO_2$ | | | | | | Aspartate feeding, 420 ppm $CO_2$ | | | | | | Pyruvate feeding | | | | | |
|---|---|---|---|---|---|---|---|---|---|---|---|---|---|---|---|---|---|---|
| | 3PGA enrichment vs $m_1$ malate | | | 3PGA enrichment vs $m_1$ aspartate | | | 3PGA enrichment vs $m_1$ malate | | | 3PGA enrichment vs $m_1$ aspartate | | | $m_2$ PEP vs $m_2$ pyruvate | | | $m_2$ PEP vs $m_2$ pyruvate | | |
| | slope | $R^2$ | *% WT* | slope | $R^2$ | *% WT* | slope | $R^2$ | *% WT* | slope | $R^2$ | *% WT* | slope | $R^2$ | *% WT* | slope | $R^2$ | *% WT* |
| | | | | | | | | | | | | | Experiment 1 | | | Experiment 2 | | |
| **WT** | 0.010 | 0.82 | | 0.011 | 0.88 | | 0.013 | 0.87 | | 0.0052 | 0.76 | | 0.016 | 0.50 | | 0.031 | 0.45 | |
| **Line 29** | 0.025 | 0.89 | *243* | 0.027 | 0.84 | *250* | 0.028 | 0.93 | *215* | 0.0084 | 0.77 | *165* | 0.043 | 0.74 | *269* | | | |
| **$ME_{BS}$ x 29-A-6-3** | 0.024 | 0.87 | *233* | 0.032 | 0.87 | *296* | 0.037 | 0.95 | *285* | 0.0092 | 0.72 | *180* | | | | | | |
| **$ME_{BS}$ x 29-C-1-5** | 0.019 | 0.93 | *184* | 0.031 | 0.94 | *287* | 0.028 | 0.87 | *215* | 0.0086 | 0.81 | *169* | 0.028 | 0.5 | *175* | 0.043 | 0.72 | *139* |
| **Line B6** | 0.029 | 0.84 | *282* | 0.046 | 0.95 | *426* | 0.054 | 0.97 | *415* | 0.0091 | 0.75 | *178* | | | | | | |
| **$ME_{BS\&M}$ x B6-B** | 0.033 | 0.75 | *320* | 0.029 | 0.84 | *269* | 0.069 | 0.90 | *531* | 0.0089 | 0.76 | *175* | | | | | | |
| **$ME_{BS\&M}$ x B6-A-5** | 0.021 | 0.71 | *204* | 0.045 | 0.90 | *417* | 0.085 | 0.90 | *654* | 0.0076 | 0.91 | *149* | | | | 0.081 | 0.9 | *261* |

**Supplementary Table 2. Data analysis to decrease experimental noise by utilizing co-variation bet-ween labelling of metabolites to estimate relative flux.** The table summarizes the slopes and $R^2$ of linear regression plots of 3PGA enrichment against fractional abundance of $m_1$ malate or $m_1$ aspartate in the [4-$^{13}$C]malate- and [4-$^{13}$C]aspartate-feeding experiments, and of the fractional abundance of $m_2$ PEP against fractional abundance of $m_2$ pyruvate in the [2,3-$^{13}C_2$]pyruvate-feeding experiment. Plots were made for wild-type, the parental lines 29 and B6 and the $ME_{BS}$ x 29 and $ME_{BS\&M}$ x B6 lines. For a given treatment and (for the [4-$^{13}$C]malate- and [4-$^{13}$C]aspartate-feeding experiments, a given $C_4$-acid $m_1$ isotopologue) the slopes indicate relative fluxes in different genotypes. Slopes cannot be directly compared across treatments and $C_4$ acid $m_1$ isotopologues because they may be differently affected by label dilution of the substrate for decarboxylation, due to compartmentation and non-steady state labelling kinetics.

In the columns '%WT', the slope in each transgenic line is given (in italics) as a percentage of that in wild-type rice. In the experiments investigating $C_4$-acid decarboxylation, compared to wild-type rice, in 420 ppm $CO_2$ the slopes for the various transgenic lines were 1.8- to 4.2-fold steeper in the [4-$^{13}$C]malate feeding experiment and 1.5- to 6.5-fold steeper in the [4-$^{13}$C]aspartate feeding experiment. In the experiments investigating PPDK activity, slopes were steeper in the transgenic lines than wild-type rice, indicating enhanced *in vivo* PPDK activity. However, as seen for fractional abundance of the PEP $m_2$ isotopologue the enhancement was smaller than for $C_4$-acid decarboxylation (compare Figure 4D with 4B-C).

The data analysis uses co-variation to cope with inherent noise in the feeding experiments. Labelled substrates were fed to single detached leaves to avoid shading by other leaves. However, any leaf-to-leaf variation in the rate of transpiration will lead to uptake of different amounts of labelled substrate. There was also considerable sample-to-sample variation in metabolite content, especially malate and aspartate (Figures S5, S6; Supplementary Datasets S1D-G). A higher endogenous content will increase dilution of incoming label. These and possibly further factors result in sample-to-sample variation in labelling intensity of the suppled substrate. Any variation in the fractional abundance of $m_1$ malate or aspartate will lead to variation in the amount of $^{13}CO_2$ that is released by decarboxylation and reassimilated into 3PGA. Similarly, variation in the fractional abundance of $m_2$ pyruvate will lead to variation in amount of $m_2$ PEP produced by PPDK. This experimental noise was decreased by comparing, on a sample-by-sample basis, 3PGA enrichment with fractional abundance of $m_1$ malate and $m_1$ aspartate in the [4-$^{13}$C]malate- and [4-$^{13}$C]aspartate-feeding experiment, and fractional abundance of $m_2$ PEP with fractional abundance of $m_2$ pyruvate in the [2,3-$^{13}C_2$]pyruvate-feeding experiment. To do this, we made scatter plots, using all individual samples at all time points (for details and plots, see Supplementary Calculations). In the malate and aspartate feeding experiments, we included $m_1$ aspartate because, the fractional abundance of $m_1$ malate may be decreased by large compartmented pools of malate that are not directly involved in metabolism [20,24-26]. The measured overall fractional abundance of $m_1$ malate may therefore underestimate fractional abundance in the malate pool that is the substrate for decarboxylation. Aspartate is not so strongly compartmented and is closely linked to metabolic malate pools via malate dehydrogenase and aspartate aminotransferase. It is plausible that $m_1$ aspartate could provide a better proxy than $m_1$ malate for the labelling intensity in the malate pool that is the substrate for decarboxylation.
